# Tracking the contribution of inductive bias to individualized internal models

**DOI:** 10.1101/2020.06.22.163295

**Authors:** Balázs Török, Dávid G. Nagy, Mariann M. Kiss, Karolina Janacsek, Dezső Németh, Gergő Orbán

## Abstract

Internal models capture the regularities of the environment and are central to understanding how humans adapt to environmental statistics. In general, the correct internal model is unknown to observers, instead approximate and transient ones are recruited. However, experimenters assume an ideal observer model, which captures stimulus structure but ignores the diverging hypotheses that humans form during learning. We combine non-parametric Bayesian methods and probabilistic programming to infer rich and dynamic individualised internal models from response times in an implicit visuomotor sequence learning task. We identify two contributors to the internal model: the ideal observer model and a Markov model capturing only immediate temporal dependencies between observations. Individual learning curves revealed internal models initially dominated by the Markov model, which was later traded-off with the ideal observer model. Thus, our results reveal a structured inductive bias that varies across individuals both in strength and persistence but is consistent in overall structure.

## Introduction

Building internal models of the environment has been a key concept for understanding human inferences in complex settings. In particular, Bayesian models of cognition provide a normative framework for understanding how humans can perform complex computations flexibly. Such models formulate the problem of learning as adaptation to the statistics of the environment. Critical tests of this concept focus on demonstrating the adaptive nature of the internal model by identifying signatures that reflect the properties of natural statistics (***Weiss et al., 2002***; ***Girshick et al., 2011***; ***Sun and Perona, 1998***) or alternatively by showing how an internal model can be established under artificial settings when conditions are precisely controlled lab settings (***Körding and Wolpert, 2004***; ***Ernst and Banks, 2002***). A particular computational appeal of Bayesian models is that these learn efficiently from limited data, and their internal representations can be flexibly used when goals are changing (***Xu and Tenenbaum, 2007***; ***Griffiths and Tenenbaum, 2009***; ***Kemp and Tenenbaum, 2009***; ***Lake et al., 2015***). However, current Bayesian models of cognition operate under a critical condition: domain-specific expert knowledge is exploited when testing the viability of the models (***Niv, 2019***; ***Maheu et al., 2020***). Here, the experimenter has to define an ideal observer model, which relates the behaviour of an observer to the process by which observations were generated. Besides the observed variables that identify the relevant features of observations, latent variables are defined, which represent quantities that are not directly available to observers but need to be inferred for efficient performance. Importantly, this ideal observer model is assumed to be readily available to humans partaking in the experiments. In simpler cases ideal observer models assess computations with low-dimensional observations and equally low-dimensional latent variables that represent natural or lab-defined features (***Körding and Wolpert, 2004***; ***Ernst and Banks, 2002***). In more complex settings internal models can also comprise scientific knowledge about the environment and are capable of supporting a rich set of possible tasks such that they can account for patterns in a wide variety of *inferences* (***Battaglia et al., 2013***). While the adaptive nature of internal models provides strong support for formulating the internal models as ideal observer models, performance in a task varies considerably across individuals. Across-individual variance motivates a more general approach that is able to identify subjective internal models based on varying behavioral responses.

Characterizing the deviation of the internal model from the ideal observer model is critical in complex environments where the exact underlying mechanism is certain to differ from the internal model maintained by human observers. This is particularly true when the observer is exposed to new environmental statistics and the model is gradually built up as information is acquired (***Love et al., 2004***; ***Gershman and Niv, 2013***; ***Nagy et al., 2018***; ***Berniker et al., 2010***). Potential sources of deviation of human performance from the one predicted by ideal observer models has recently attracted intense attention (***Rahnev and Denison, 2018***; ***Song et al., 2019***). Studies have demonstrated that learning a novel and complex statistics can lead to systematic deviations from the ground truth model (***Roach et al., 2017***; ***Gekas et al., 2013***). Mismatch between the predictions of an ideal observer model and human behaviour has been shown to be a consequence of computations relying on an internal model that deviates from the ground truth rather than suboptimal computations (***Acerbi et al., 2014***; ***Drugowitsch et al., 2016***; ***da Silva and Hare, 2020***). Acquiring the right internal model during learning is a daunting task, which in general cannot be efficiently accomplished based on the actual observations alone. These results point to an important insight: when learning about a new environment, however simple it might be, humans not only rely on actual experiences but recruit internal models that they can harness in this new environment. Relying on previous experience by recruiting previously acquired internal models is a rational strategy when available data about a particular environment and/or task is limited but has the consequence of potentially inducing biases in the interpretation of new data (***Griffiths et al., 2010a***). Indeed, such inductive biases are readily recruited when interpreting the mechanics during the manipulation of virtual objects (***Neupärtl et al., 2020***). Taken together, inductive biases and a model that summarizes our knowledge about the environmental statistics jointly determine our responses. Therefore we aim to track the deviation of the internal model from the ideal observer models to disentangle the contributions of the internal model and the inductive biases to the behavioral responses and ultimately to uncover inductive biases.

To capture the internal model that underlies an adaptive behavior but can be richer than the ideal observer model and can reflect the differences between individuals arising from subject-specific inductive biases and different learning paths, we need both a (i) highly expressive class of internal models and (ii) behavioural measurements that are highly informative about the internal model. We proceed by choosing an experimental paradigm that satisfies (ii). In an experiment where trials are governed by temporal dynamics and therefore individual trials are not independent, the joint set of behavioural measures in a number of trials have information content that far exceeds that of an independent and identically distributed (i.i.d.) experimental setting. In this paper, our goal is to recover dynamical internal models of individuals in a dynamic task novel to participants. It has been extensively documented that participants do pick up temporal regularities in experiments with stochastic dynamics (***Glaze et al., 2015***; ***Urai et al., 2017***; ***Braun et al., 2018***). Furthermore, individuals show high variation in their initial assumptions (***Glaze et al., 2015***; ***Mathys et al., 2014***). In order to satisfy (i), we propose to use infinite Hidden Markov Models (iHMMs, ***Gael et al. 2008***) as a model class to characterize the internal models maintained by individuals. To infer the structure and dynamics of the iHMM we adopt and extend the Cognitive Tomography framework (***Houlsby et al., 2013***). Cognitive Tomography (CT) aims to reconstruct high-dimensional latent structure using low-dimensional behavioural measurements. Critically, CT distinguishes two components underlying behavioral responses: (1) the internal model which captures the statistical regularities of the environment, independent of the behavioral responses associated to a particular task (task-general component), and (2) a response model that links the individual’s inferences to their observable behaviour (task-specific component). In its original formulation the internal model structure was assumed to be unaffected by the observations during the experiment, the internal model was not dynamical, and behaviour was characterized solely by the key presses participants performed essentially providing roughly one bit of information in each trial. According to our model, participants filter the information gained from their observations over time, combine this information with their estimate of the dynamics to predict the next stimulus. We relate their subjective predictions to response times using the linear ascend to threshold with ergodic rate (LATER) model (***Carpenter and Williams, 1995***). We invert this generative model of behaviour to infer individuals’ internal models.

In order to investigate the characteristics of internal models discovered by individuals when exposed to a novel stimulus sequence, we developed CT to operate on dynamical datasets and response time data. Specifically, using CT we wanted to have an insight into the type of inductive biases humans might have and into the ways they balance these with evidence coming from observations. We use an implicit learning paradigm in which the stimulus sequence is characterised by a challenging statistics novel to participants and track behavioral responses over the course of multiple days. We hypothesise that the dynamical properties of the natural environment imply strong expectations for sequential dependencies between subsequent stimuli, known as Markovian structure, therefore we explore the contribution of such a Markovian inductive bias to the internal model of individuals. To achieve this, the stimulus statistics used in the experiment lacks a Markovian component and is solely characterised by higher-order temporal dependencies. We infer individual learning curves using CT to have an insight into the varying inductive biases participants maintain and also into the diverging level of integration of evidence with initial biases. First, we develop CT to provide participant-by-participant and trial-by-trial prediction of response time data and demonstrate that CT outperforms alternative models, most notably the ideal observer model of the task, which relates behavioural responses to the ground truth model of stimulus generation. Second, to demonstrate that the internal model found by CT is indeed representing the belief of a participant about the statistical structure of stimuli, we perform a pair of analyses to demonstrate that we can successfully dissociate the internal model and the behavioral model. Third, we show that even after eight days of training there are substantial differences between individuals and learning curves display diverging patterns. Importantly, while the contribution of the ideal observer model increases with training, the internal model discovered by CT substantially outperforms the ideal observer model in most individuals. Fourth, we show that the performance of CT can be largely explained by combining the Markov and ideal observer models, therefore the contribution of a Markovian inductive bias and evidence can be assessed on a participant-by-participant basis. Finally, we evaluate relevant internal variables of the internal model that can provide regressors for neural activity. Taken together, our results suggest a new perspective on how humans trade-off inductive biases and evidence over the course of learning and also provide new tools to measure such inductive biases.

## Results

In order to test how behavioural data from individuals can be used to infer a dynamical probabilistic latent variable internal model and assess the contribution of inductive biases to the internal model, we used an experimental paradigm that could fulfil a number of key desiderata. First, the paradigm relies on across-trial dependencies; second, as in everyday tasks, the state of the environment cannot be unambiguously determined from the observation of momentary stimuli; third, the structure of the task is new to participants; fourth, the complexity of the task is relatively high, i.e. an a priori unknown number of latent states determine the observations; fifth, behavioural measurements during task execution are continuous, which ensures that rich inferences can be made. In the alternating serial response time task (ASRT, ***Howard and Howard 1997***) a stimulus can appear at four locations of a computer screen and the sequence of locations (untold to participants) follows a pre-specified structure (1A). In odd trials, the stimulus follows a 4-element sequence, while in even trials the stimulus appears at random at any of the positions with equal probability independently of all other trials (1, SI Appendix). Such stimuli precluded unambiguously determining the state of the task solely based on a single trial’s observation. There are an additional 5 random trials at the beginning of each block. Participants are tasked to give fast and accurate manual responses through key presses corresponding to the locations of the stimuli. We collected response time measurements for sequences of stimuli organized into blocks of 85 trials. A session consisted of 25 blocks and the performance was tracked for 8 days with one session on each day, during which the same stimulus statistics governed the stimuli, followed by two additional sessions on subsequent days where the statistics of stimuli was altered (SI Appendix Fig. S1).

We used the response times of individuals to infer a dynamical probabilistic latent variable model underlying their behaviour (Fig. 1, SI Appendix). We invoked the concept of CT to infer the internal model from a limited amount of data. CT requires the formulation of the generative model of the data, i.e. the process that produces behavioral data from observations. CT distinguishes two components of the model (1B, *grey boxes*): the internal model, which summarizes an individual’s knowledge about the stimulus statistics and the behavioral model, which describes how behavioral responses are related to the internal model during the task that is being performed. Inference of the internal model requires inference how latent states evolve and how these determine the stimuli. By knowing the dynamics of latent states we can make predictions for the upcoming stimuli by establishing the predictive probability of possible subsequent elements of the stimuli. The behavioral model establishes how predictive probabilities of the internal model are related to behavioral outcome, which is the response time in our case. We used the LATER model to predict response times from predicted probabilities (***Carpenter and Williams, 1995***; ***Noorani and Carpenter, 2016***). The experimenter uses the observed data (1B, grey boxes), the stimulus sequence and response times, for the inference. The resulting CT model (SI Appendix Fig. S2) is implemented as a probabilistic program with components implemented in Stan (***Carpenter et al., 2017***).

**Figure 1.**
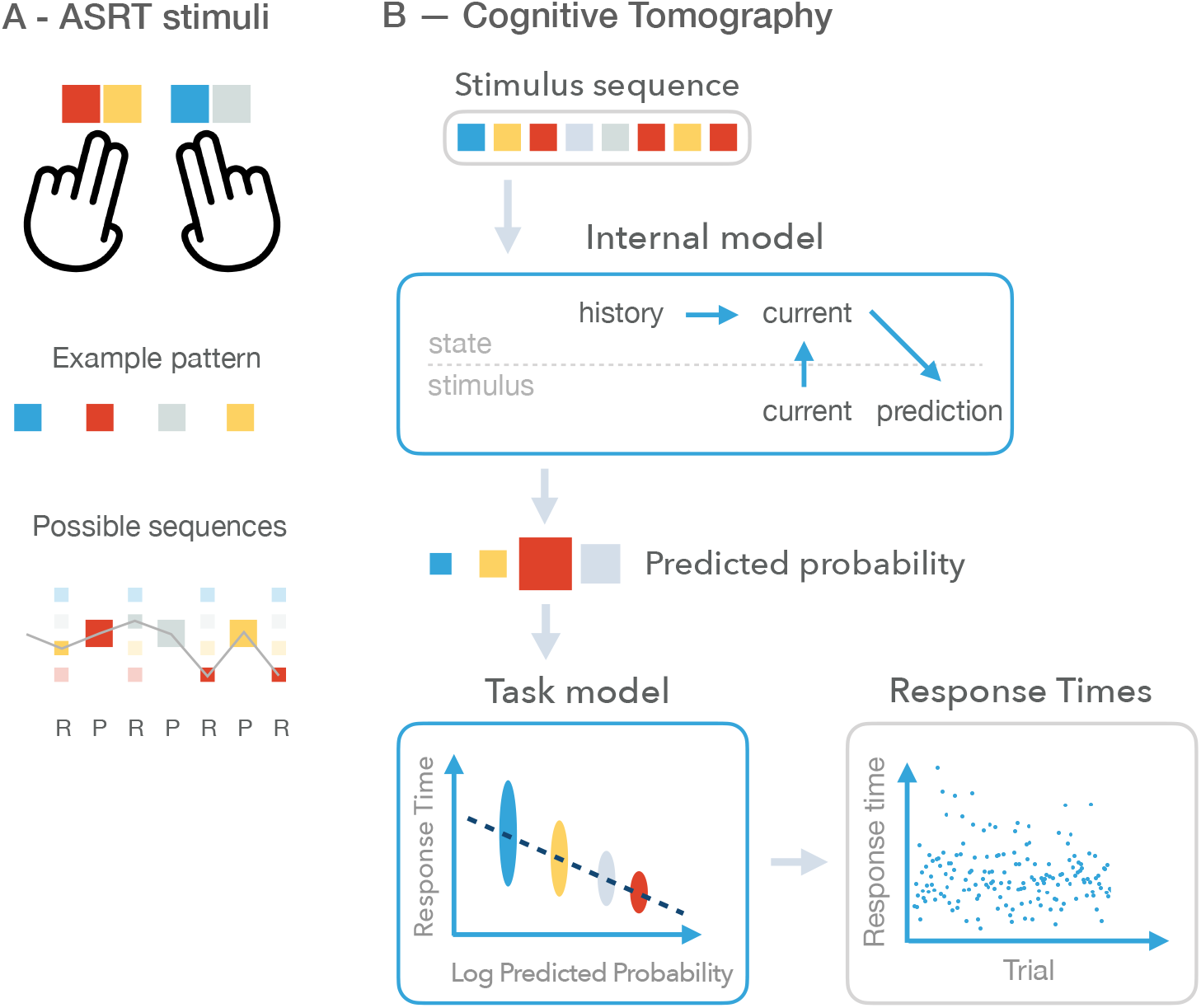
Experimental paradigm and Cognitive Tomography (CT). **A** *Top:* Behavioural responses: participants are responding with key presses on a keyboard where stimulus identities (shown as different *coloured squares*) are associated with unique keys. *Middle:* An example deterministic pattern sequence, which recurrently occurs in the stimulus sequence of a particular participant. Different participants are presented with permutations of this four-element sequence. *Bottom:* In the actual stimulus sequence presented to participants, the deterministic pattern sequence is interleaved with random items (*small squares.* Random items can be any of the four stimuli and can occur with equal probability (*size* of the square is proportional to the probability of a stimulus). *Grey line* indicates one particular realization of the stochastic sequence. **B** The probabilistic generative model underlying Cognitive tomography. The generative model describes the process how a stimulus sequence (*top grey box*) results in a behavioural response. A participant is assumed to use the internal model *top blue box* to make a prediction for the upcoming stimulus. The internal model assumes dynamics over the latent states. The current latent state is determined jointly by earlier states and the current observation. Based on the current latent state a prediction can be made on the probability of possible upcoming stimuli. The predicted probability (*size of squares* corresponds to the probability of prediction) is related to the behaviour through a behavioral model (*bottom blue box*). The behavioral model depends on the task being performed and therefore the type of response being predicted. Here, the logarithm of the predictive probability is mapped to a mean response time and actual response times are assumed to be noisy versions of this mean. Response times (*bottom grey box*) shown here are 400 trials from an example participant. Cognitive tomography uses the stimulus sequence and the sequence of behavioural responses (*grey boxes*) to infer the components of CT, the internal model and the behavioral model (*blue boxes*).

The iHMM model provides a flexible model class to infer latent variable models (***Gael et al., 2008***). Similar to the classical Hidden Markov Model, learning requires the specification of transition probabilities between latent states along with the probability distributions of observations given a particular latent state (2A). Additional flexibility of iHMM is provided by not fixing the number of latent states but inferring this from data. This is implemented as a non-parametric Bayesian model. In an iHMM, participants filter the information gained from the observations over time to estimate the possible latent state of the system (Fig. 2Ba, *filled purple circles*). That is, they infer what history of events could best explain the sequence of their stochastic observations. Then, they use their dynamical model to play the latent state forward (Fig. 2Ba, *open purple circles*) and predict the next stimulus (Fig. 2Bb).

**Figure 2.**
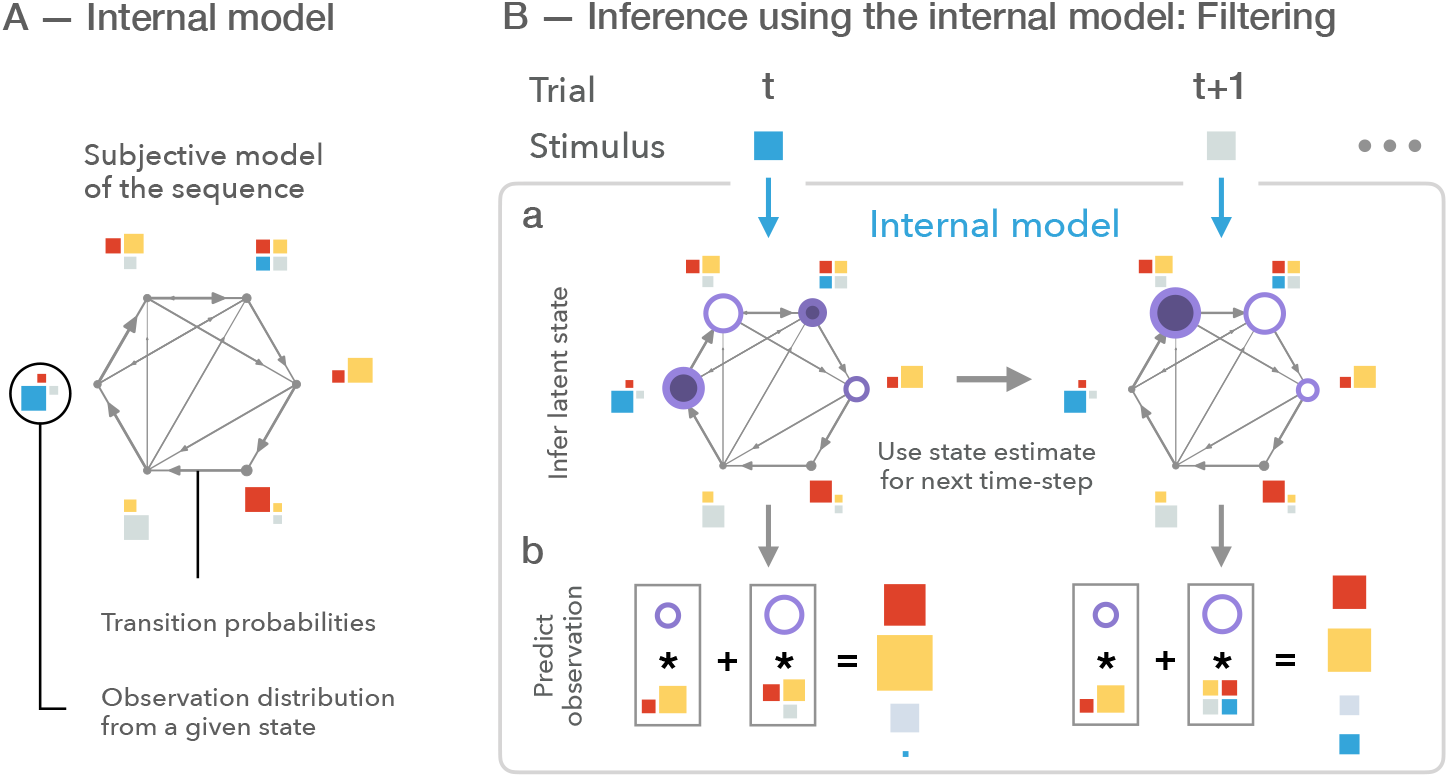
Inference and predictions using the internal model. **A** We formulate the internal model as an iHMM, where the number of latent states (*grey dots*), transitions between the states (*arrows*), and the distribution of possible stimuli for any given state (*coloured squares*) needs to be inferred by the experimenter. Width of arrows is proportional to transition probability and arrows are pruned if the transition probability is below a threshold; size of dots indicates the probability of self-transition. Size of stimuli is proportional to appearance probability in the given state. The result of inference is a distribution over possible model structures, the figure represents a single instance of such a distribution. **B** Evolving the internal model from trial *t* to trial *t* + 1. At time *t*, participants use the internal model components to update their beliefs over the current state of the latent states (**Ba**, size of *dark purple discs* represent the posterior belief of the latent state based on the current observation, *blue square*). Then, participants play the model forward into the future (*open purple circles*). Finally, they generate predictions for the upcoming stimulus (**Bb**, *squares in grey boxes*) by summing over the possible future states (*open purple circles in grey boxes*). Participants use previous state beliefs and the new stimulus to update latent state beliefs. In this particular example, at trial *t* + 1 only one of the possible states can generate the observation, hence there is only one dark purple disk. Again, they play the dynamics forward and predict the next stimulus.

To test that the proposed inference algorithm is capable of the retrieval of the probabilistic model underlying response time sequences, we validated our inference algorithm on synthetic data (SI Appendix, SI Appendix Fig. S3). We used three different model structures for validation, which were HMMs inferred from three different-length stimulus sequences (one sample from the iHMM inference in ***Gael et al. 2008***). Similar to our human experiment data, we assessed CT by computing its performance on synthetic response times. Further, since synthetic participants provide access to true subjective probabilities we also calculated performance on the ground truth subjective probabilities. We showed that the subjective probabilities can be accurately recovered from response times. As shown on SI Appendix Fig. S3C, standard deviations of participants’ response times are within the range of successful model recovery.

### Alternative models

The combination of a non-parametric Bayesian model and probabilistic programming provides the CT model with the flexibility to accommodate a wide spectrum of possible internal models that ranges from very simple internal models relying only on a very limited number of latent variables or substantially more complex corresponding to higher number of latent variables. In order to understand the characteristics of the internal model we need to have analysis methods that provide us with the necessary insights. To do so, we sought to decompose the internal model into components that can reveal the computational principles governing it.

Whether and how much the inferred internal model reflects the structure of the environment can be tested by contrasting the inferred CT model with the ideal observer model. Since we have full control over the generating process of the observed stimuli, the ideal observer model is identified with a generative model that has complete knowledge about the stimulus statistics and the only form of uncertainty afflicting inference stems from the ambiguity in the interpretation of observations rather than uncertainty in model structure or parameters. Since the structure of the task was fixed, this model could not account for possible across-individual differences in internal models. Assessment of the deviation of the CT and the ideal observer models can reveal the richness of the strategies pursued by humans when exposed to unfamiliar artificial stimulus statistics. Fixed parameters of the ideal observer also ensured that the changing performance of humans could be directly compared across the course of learning to the same baseline. Importantly, ‘learning’ by participants during extended exposure throughout the experiment does not necessarily mean that their internal model gets gradually closer to the ideal observer model since even when more evidence is provided towards the true underlying model one can commit more and more to a superstitious model. As a consequence, deviation can temporarily accumulate before converging towards the true model, resulting in nonlinear learning trajectories.

A defining characteristic of the ideal observer model was that it assumed latent states underlying observations. An internal model that lacks this critical assumption can model how such a false but feasible assumption might be able to predict response times. We used a Markov model to implement this assumption, which can model dynamics but can only represent transitions directly between observations without the introduction of latent variables. Such a model can capture individual differences and, importantly, a Markov-like dynamics is a special case of the internals models that CT can represent.

Finally, the gold-standard for characterizing learning in an ASRT task is the so-called triplet model that tests the correlation of response times with the summary statistics of the stimulus sequence and we formulated as a trigram model. We include this model as well to compare its performance with alternative models. Since the triplet model reflects the summary statistics of the stimulus this model bears resemblance to the ideal observer model albeit without assuming the ability to perform real inference in the model.

Direct comparison of the alternative models is summarized in Table 1 (see also SI Appendix).

**Table 1.** Table of models and the maximum likelihood parameter sets for the stimuli in our experiment. The ideal observer model (the true generative model of the stimuli) can be formalized as an 8-state HMM with states Pattern1, Random1, Pattern2, Random2, Pattern3, Random3, Pattern4, Random4 where the pattern states produce the corresponding sequence element with probability 1 and all the random states produce any of the four observations with equal probability independently. The Markov model (where predictions are produced by conditioning only on the previous observation) fits the observations best when it predicts all observations with equal probability, since the marginal probabilities of any one stimulus is equal regardless what the previous observation was, because every other trial is random. The trigram model produces a “high triplet” prediction, where the next stimulus is the successor of the stimulus two trials ago in the pattern sequence. All alternatives have equal probability of 0.125. Note that the exact probabilities in this case are not relevant since the trials are categorized into two groups (high and low) and therefore the parameters of the response time model and these probabilities are underspecified. The CT model produces a prediction for the next stimulus via filtering. A latent state of the sequence is estimated from previous observations using a Hidden Markov Model. This flexible model space includes the ideal observer model as well as the Markov model as special cases.

| Model space |  | Best model<br>in model space |
| --- | --- | --- |
| Ideal Observer | $p(y_{t+1}\vert y_1, y_2, \dots, y_t)$ | 8-state HMM<br>deterministic latent dynamics<br>stochastic observations |
| Markov | $p(y_{t+1}\vert y_t)$ | Uniform |
| Trigram | $p(y_{t+1}\vert y_t, y_{t-1})$ | (0.625, 0.125, 0.125, 0.125)<br>"high" & "low" |
| CT | $\sum_{s_t, s_{t+1}} p(y_{t+1}\vert s_{t+1}) \cdot p(s_{t+1}\vert s_t) \cdot p(s_t\vert y_1, y_2, \dots, y_t)$ | Ideal Observer |

### Trial-by-trial prediction of response times for trained participants

We inferred the internal model along with the parameters of the response time model on 10 blocks of trials measured at the second half of the session. Similarly, all the alternative models were fitted on the same data. To demonstrate that all the models are capable of predicting the empirical response time distribution, we synthesized response time data from all four models (Fig. 3A). Distribution of synthetic response time distributions matched the empirical distribution closely (Fig. 3B). The response time model assumes that variance in response times comes from the joint effect of the variance in log predictive probabilities and a Gaussian noise corrupting the subjective probabilities. If the fit of the internal and the response models are appropriate, the residual variance not accounted for by the subjective probabilities is expected to be normally distributed. We checked this on the CT model on a subject by subject basis, which analysis demonstrated that residuals are close to a normal distribution (SI Appendix Fig. S4) with a single subject apparently having a bimodal residual distribution, potentially indicating additional structure in the internal model not captured by CT. Note, that throughout the analysis trials with fast response times are discarded (see Methods for details).

**Figure 3.**
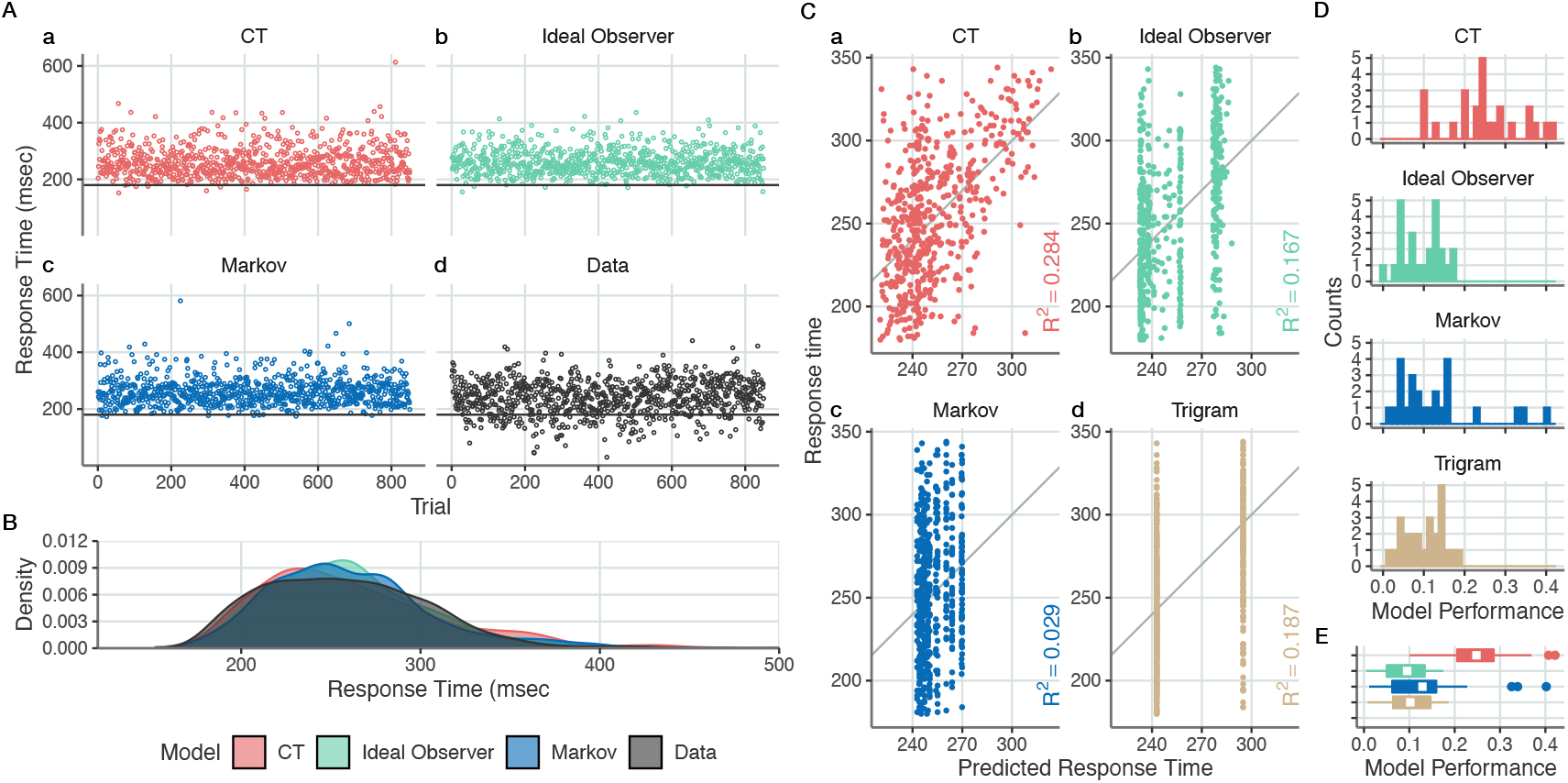
Response time predictions for individual trials on a held-out test dataset. After training our inference algorithm on a training dataset of 10 blocks, we predict response times of another 10 blocks on the same day by the same participant. **A**, Synthetic response time data for three models (CT, **Aa**; ideal observer model, **Ab**; Markov model, **Ac**) in modelled trials and the empirical data from an example participant (**Ad**). **B** Marginal distributions for the different models and for the empirical data. **C** Individual trial predictions against measured response times (*dots*) on held-out data for an example individual by all the four models considered. Response times shown between 180msec and mean+3 s.d. for visual clarity. Performance is measured as the trial-by-trial correlation between measured and predicted response times (*R*^2^, *coloured labels*). **D** Trial-by-trial model performance histograms (*R*^2^ as a measure of explained variance) of the different models for all participants on Day 8 of the experiment. **E** Summary of trial-by-trial model performances (*R*^2^ values) of the experiment for all participants on day 8. White rectangles show means, hinges of boxplot show 25th and 75th percentiles. Upper whiskers show largest value or 75th percentile + 1.5 · *IQR.* Similarly for lower whisker. Color code as on panels **C** and **D**.

A more relevant test of the models concerns the prediction of response times on held-out data. We tested the performance of all models on 10 blocks of trials from the first half of a session. Model performance was measured by evaluating the amount of variance of response times explained (*R*^2^) on the held-out response times in individual trials (Fig. 3C), which provides a levelled comparison for the Bayesian and non-Bayesian accounts. In our predictions we show the maximum-probability response times resulting in inhomogeneous bands in the predicted distributions. Response times could be predicted by CT efficiently even for individual trials as shown by the analysis of the response times from a single participant *R*^2^(550) = 0.284, *p* < 0.001, Fig. 3Ca). It is important to note that the predictive power was substantially increased by averaging over trials in the same positions of the sequence (SI Appendix Fig. S5). Despite the significant advantage of trial-averaged predictions, we believe that single trial predictions provide a more rigorous and important characterization of human behaviour therefore we evaluate model performances on an individual trial basis in the rest of the paper. Predictive power of CT varied across participants but correlations between measured and predicted response times was significantly above zero for all participants (*M* = 0.248 ranging from 0.1 to 0.42, Fig. 3Ca).

Assessment of all the models on individual subjects enables us to directly compare the relative predictive power of the three models (Fig. 3C). Note, that plotting the maximum probability prediction results in predictions that are more stratified than the empirical data despite having smooth predictive distributions (Fig. 3B). For the example participant analysed here, the ideal observer model had a significantly lower performance than the CT model, indicating that CT captures more structure in the response times (Fig. 3Cb, binomial test on MSE values 0.96, *n* = 25, *p* < 0.001), potentially indicating a more accurate internal model than the ideal observer model. Outperforming the ideal observer might look surprising but the ideal observer assumes that the internal model is solely and completely determined by the stimulus statistics, which is not necessarily true for the internal model entertained by human observers. The performance of the Markov model was substantially lower, still distinct from zero (Fig. 3Cc, *R*^2^(550) = 0.029, *p* < 0.001). The trigram model showed a lower performance than CT, with a performance similar to that of the ideal observer. All models could predict response times on a trial-by-trial basis with significantly above zero performance (Fig. 3D), but on average controls had substantially lower performance than that of CT (CT vs Trigram 0.92, *n* = 25, *p* < 0.001 CT vs Markov 1, *n* = 25, *p* < 0.001 CT vs Ground Truth 0.96, *n* = 25, *p* < 0.001, Fig. 3E).

### Validation of the internal model

To verify that better predictive performance of the model identified by CT is not only a consequence of a more flexible model but is a signature of inferring a better internal model, we perform two additional analyses. The core principle of Cognitive Tomography is to distinguish an internal model that captures a task-independent understanding of the statistical structure of the environment and a response model which describes task-specific behavioural responses. In order to assess the contribution of inductive biases and to assess how evidence is integrated with inductive biases, we need to validate that the internal model found by CT is indeed reflecting the belief individuals have on the statistical structure of stimuli and not the task being executed. This validity therefore hinges upon the validity of the task-dependent behavioral model, which can be assessed by manipulating the internal and behavioral models independently. First, we tested if the same internal model can be used to predict behavioral performance in a different task. We consider a task different if a new behavioral measure is predicted even though the participants are engaged in the same psychophysics task, the ASRT in our case. So far, we predicted response times when participants pressed the correct key. We replaced this task with the prediction of behavior in error trials. In particular, we aimed at predicting the trials in which a participant is likely to commit errors and also the erroneous response when an error occurs. Second, we tested the usage of the internal model when stimulus statistics is manipulated. After completing eight days of training, participants were exposed to a novel stimulus statistics and we tested if participants recruited the learned internal model only when the stimulus statistics matched the one the internal model had been learned on.

In error prediction we separated trials based on whether the participant pressed the key corresponding to the actual stimulus or any other keys. Note, that the internal models of CT were inferred only on correct trials using the response time model. We investigated two relevant hypotheses. First, a participant will more likely commit an error when their subjective probability of the stimulus is low. Second, when committing an error, their response will be biased towards their expectations. For reference, we contrasted the performance of CT both with the ideal observer model and the Markov model. We compared the rank of the subjective probability of the upcoming stimulus both for correct and incorrect trials (Fig. 4A). CT ranked highest the upcoming stimulus in correct trials above chance (0.461, *n* = 18473, *p* < 0.001) and significantly below chance for incorrect trials (0.175, *n* = 2777, *p* < 0.001). Ideal observer model excelled at predicting the correct responses, as it ranked the correct responses high above chance (0.635, *n* = 18473, *p* < 0.001). However, it also assigned the highest probability to the upcoming stimulus in incorrect trials (0.315, *n* = 2777, *p* = 1). The Markov model’s performance was close to chance in predicting the upcoming stimulus in correct trials (0.271, *n* = 18473, *p* < 0.001). In incorrect trials the Markov model, too, showed a tendency to not rank the upcoming stimulus the highest, therefore showed significantly below chance performance on ranking (0.137, *n* = 2777, *p* < 0.001). Ranking of incorrect responses was above chance for all models (Fig. 4B).

**Figure 4.**
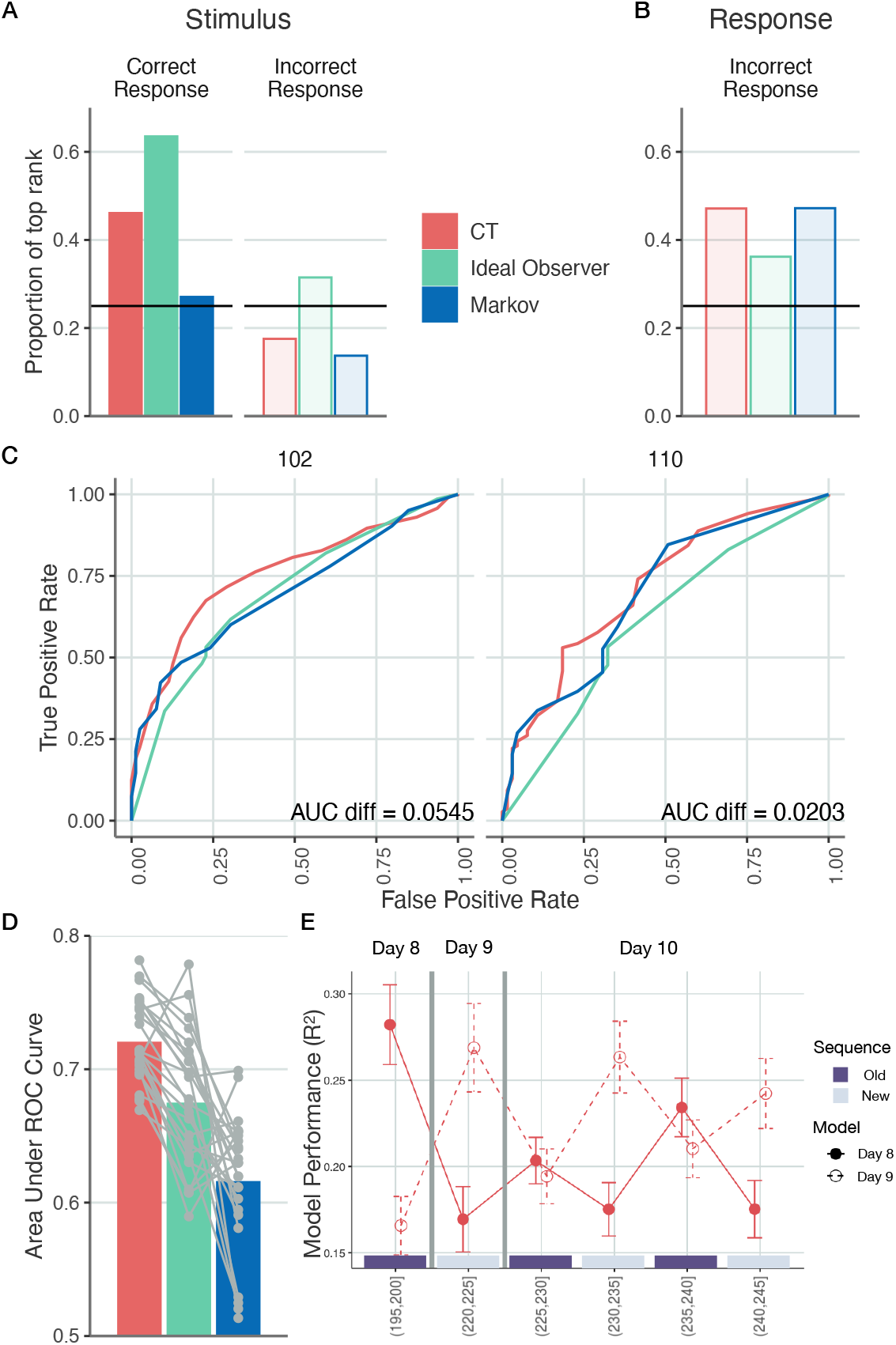
Validation of the inferred internal model by selectively changing the task and the stimulus statistics. **A-D** Choice predictions by CT (*red*), ideal observer (*green*), and Markov models (*blue*). Models are trained on response times for correct key presses on Day 8 and tested on both correct and error trials the same day. **A**, Proportion of trials where the model ranked the upcoming stimulus first. For correct trials, the Markov model has only marginal preference for the stimulus. For incorrect trials, the ideal observer model falsely predicts the stimulus in more than a quarter of the time. **B**, Proportion of trials where the model ranked the button pressed by the participant first. For incorrect responses, all models display a preference towards the actually pressed key over alternatives. **C**, ROC curves for two example participants based on the subjective probabilities of upcoming stimuli (held-out dataset). Area under the ROC curve characterizes the performance of a particular model in predicting error trials. **D**, Area under ROC curve. Grey dots show individuals, bars show means. **E** Investigating new internal models emerging as new stimulus sequences are presented. Performance of predicting response times on Day 8-10 using CT-inferred models that were trained on Day 8 (*filled red symbols*) and Day 9 (*open red symbols*) on stimulus sequences governed by Day 8 or Day 9 statistics. On Day 9 a new stimulus sequence was introduced, therefore across-day prediction of response times corresponded to across sequence predictions. Training of the models was performed on 10 blocks of trials starting from the 11th block and prediction was performed on the last five blocks of trials (the index of the blocks used in testing is indicated in brackets). On Day 10, stimulus sequence was switched in 5-block segments between sequences used during Day 8 and Day 9 (*purple and grey bars* indicate the identity of stimulus sequence with colours matching the bars used in Day 8 and Day 9. Error bars show 2 s.e.m. Stars denote *p* < 0.05 difference.

We obtained a participant-by-participant assessment of the difference between model performances in predicting error trials by calculating ROC curves of the models based on the subjective probabilities assigned to upcoming stimuli (Fig. 4C, SI Appendix Fig. S6). Area between two ROC curves characterizes the performance difference between models and CT is shown to consistently outperform both the ideal observer and Markov models in distinguishing correct choices from incorrect choices (paired t-test on AUC values one-sided *t(24) =* 6.185, *p* < 0.001, *CI* = [0.033, *Inf*], *d* = 1.1 paired t-test on AUC values one-sided *t*(24) = 9.584, *p* < 0.001, *CI* = [0.0858, *Inf*], *d* = 2.48 for ideal observer and Markov models, respectively, Fig. 4D). Thus, the ideal observer model is substantially outperformed by CT in across-task predictions and the Markov model was shown to outperform the ideal observer model in predicting incorrect responses.

In order to test how participants use and update their internal model when stimuli are generated using a new stimulus statistics, participants were tested in two additional days. On Day 9 of the experiment, 25 blocks of trials were performed with a new pattern sequence. On Day 10, another 20 blocks of trials were performed switching between the pattern sequences of Day 8 and Day 9 every five blocks. Comparison of within-day and across-day predictions on Days 8 and 9 demonstrates that response time statistics change across days, as reflected by significant differences in within-day and across-day predictions (one-sided *t*(24) = 4.958, *p* < 0.001, *CI* = [0.0746, *Inf*], *d* = 1.06 and one-sided *t*(24) = 4.9, *p* < 0.001, *CI* = [0.0616, *Inf*], *d* = 0.963 for forward and backward predictions, respectively). Specificity of response time statistics to the stimulus statistics is tested by predicting Day 10 performance using Day 8 and Day 9 models. The Day 8 model more successfully predicts response times in blocks relying on Day 8 statistics than on blocks with Day 9 statistics (one-sided *t*(24) = 3.734, *p* < 0.001, *CI* = [0.0236, *Inf*], *d* = 0.594) and the opposite is true for the Day 9 model (one-sided *t*(24) = 3.528, *p* < 0.001, *CI* = [0.0261, *Inf*], *d* = 0.575). Oscillating pattern in the predictive power of Day 8 and Day 9 models on blocks governed by Day 8 and Day 9 statistics indicates that participants successfully recruit different previously learned models for different stimulus statistics (Fig. 4E and SI Appendix Fig. S7).

### Evolution of the internal model with increased exposure

In order to assess how an initially assumed internal model might give space to an internal model that reflects the statistics of the stimuli, we tracked the evolution of the internal models during eight days of training using the same stimulus statistics. Learning the model underlying observations entails that participants need to learn the number of states, the dynamics, and observation distributions. As a consequence, learning is challenging in the ASRT paradigm and therefore requires substantial exposure to stimulus statistics. We established individual learning curves by learning separate models for different days (Fig. 5A). Note, that this approach can identify changes in the internal model on the time scale of hundreds of trials (corresponding to different days in our paradigm) but assumes constancy of the internal model within this particular number of trials. To have an insight into how the statistics captured by CT changes over the days of training, we compared its performance to the ideal observer model. Besides the ideal observer model that cannot capture individual differences, we also included the Markov model in the analysis. While this model only captures immediate dependencies between trials, statistics simpler than the true statistics underlying stimuli, it has the benefit that it can capture individual differences. Finally, we also included the trigram model in this analysis, which can capture essential summary statistics of the stimuli. The statistical structure captured by the ideal observer is only weakly present after a day of training (one-sided *t*(24) = −0.7692, *p* = 0.775, *CI* = [-0.0214, *Inf*], *d* = 0.154, Fig. 5A) but the Markov model can capture a significant amount of variance from response times (*M* = 0.0677 ranging from 0.00049 to 0.13, one-sided *t*(24) = 11.68, *p* < 0.001, *CI* = [0.205, *Inf*], *d* = 2.34), and it is not different from CT (binomial test on MSE values 0.72, *n* = 25, *p* = 0.0433). Since CT contains the Markov model as a special case, this result suggests that after a single day of training it is a simple Markov structure that dominates the responses of participants. The performance of the trigram model closely follows that of the ideal observer, indicating that the summary statistics captured by the trigram model is indeed responsible for a substantial part of the statistics reflected by the ideal observer.

**Figure 5.**
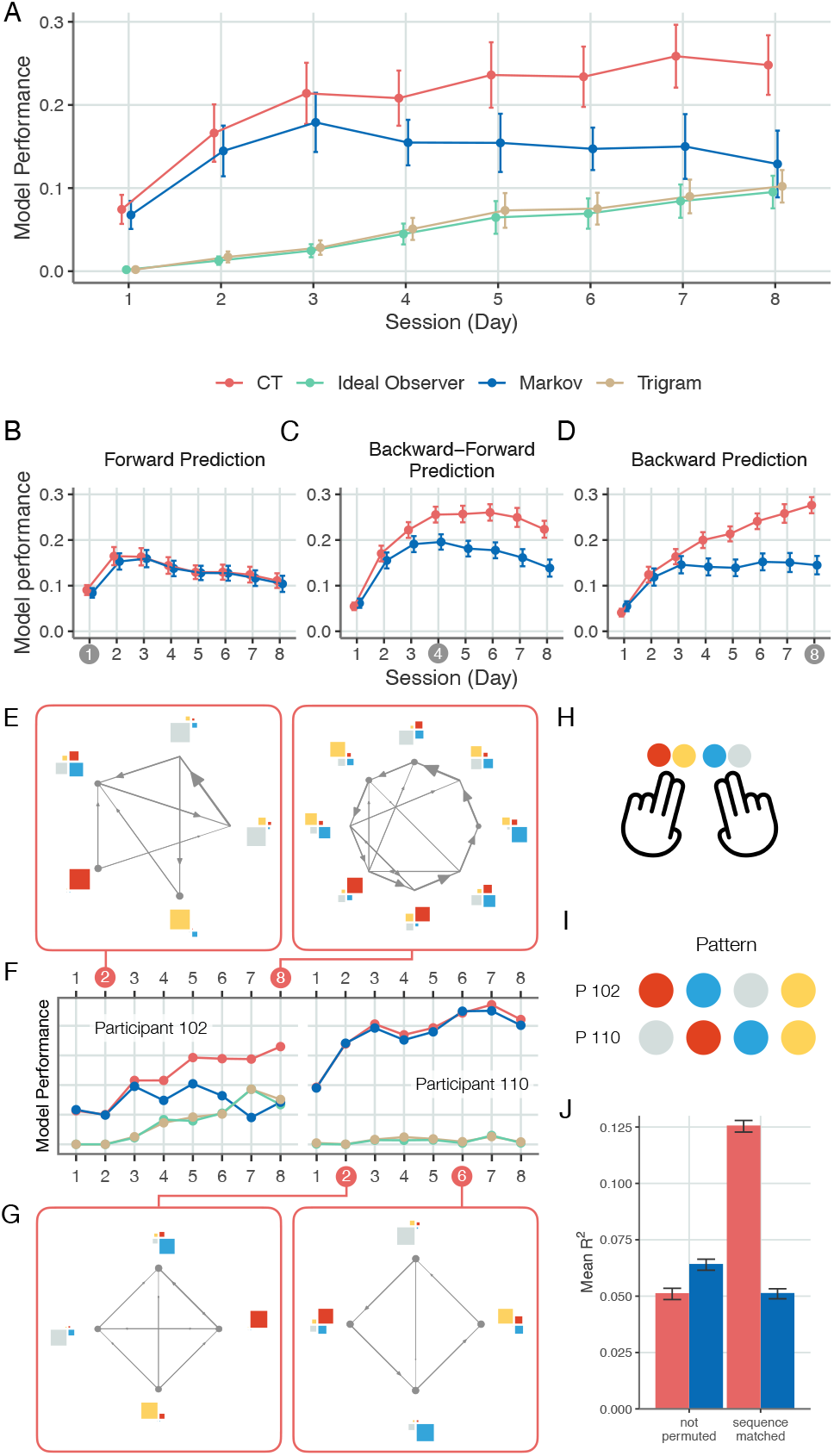
Evolution of the internal model with increasing training. **A** Mean explained variance (*dots*, averaged over participants) in held-out response times in sessions recorded on successive days for the CT (*red*), Markov (*blue*), ideal observer (*green*) and trigram (*yellow*) models. Error bars denote 2 standard error of the group mean. **B** Forward prediction performance of the CT and Markov models across different days of the experiment. Generalization capacity of models were tested by training models on the first day of experiment and testing them on successive days. We inferred internal models on the first day of the experiment and used this internal model to predict further days’ response times of the same participant. Dots show means of model performances (*R*^2^ values) over participants. Performance of the training day is measured on held-out trials. **C** Forward-backward prediction of CT and Markov models. Same methodology is used as on panel **B** but for models trained on Day 4 and tested both on preceding and succeeding days of the experiment. **D** Backward prediction of CT and Markov models. Model training was performed on Day 8 and tested on preceding days. **E-G** Learning curves of individual participants characterized by different models. **E** Participant 102 finds a partially accurate model by Day 2 and a model close to the true model by Day 8. **F** Individual learning curves for two example participants (102 and 110). Colour code same as **A**. **G** Participant 110, sticks to a Markov model and prediction of their behaviour by this model is gradually improves. This is further supported by the ideal observer model’s floored performance, indicating that no higher-order statistical structure was learned. **H** Color coding of response buttons used in this figure. **I** Color coding of sequence showed to participants. **J** Prediction of response times across participants. Day 8 internal models are swapped across participants and performance of swapped models is measured (*left*) both for CT (*red*) and the Markov model (*blue*). We construct sequence-matched models by inverting the permutation (*right*). Error bars show 2 s.e.m.

Performance of all models increases in the first few days of the experiment but the advantage of CT gradually increases as time passes (*F*(7,168) = 32.43, *p* < 0.001, 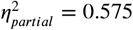) and by day eight the dominance of CT model is very pronounced, reaching *R*^2^ = 0.248 (s.e.m. 0.0179) on average.

By using a model trained on a specific day to predict responses on subsequent days (forward prediction) or previous days (backward prediction), we can obtain insights into the dynamics of learning the internal model. Participant-averaged backward prediction in CT indicates a gradual build-up of the model as prediction of earlier day responses strongly depended on the time difference between the training day of the model and the test day (Fig. 5C,D). Forward prediction did not exhibit such strong dependence on time difference, indicating that on group level, the statistical structure acquired up to a particular day was incorporated in later models (Fig. 5B,C). Importantly, group averages can hide differences between individual strategies but CT provides an opportunity to investigate these through the patterns of forward an backward predictions (SI Appendix Fig. S8). For instance, participant 119 initially entertains a Markov-like internal model up until day 2-3, only to dispose of it and acquire a different structure later in the training, indicating that the discovery of a more complex statistical structure can completely eliminate the contribution of the Markov model to behavioural responses.

CT offers a tool to investigate the specific structure of the model governing behaviour at different points during learning (Fig. 5E,G). We computed individual learning curves (Fig. 5F & SI Appendix Fig. S9) by measuring the performance of the different models on each day of the experiment. Furthermore, we analyse the internal model structure associated with behaviour by taking posterior samples from the CT model. Early during training where the performance of the Markov model is close to that of CT, the inferred iHMM indeed has a structure close to that of the Markov model, which is characterized by a strong correspondence between observations and states. Later in the experiment, however, the performance of CT deviates from that of the Markov model for most of the participants (SI Appendix Fig. S9) and the model underlying the responses is close to the ideal observer model (Fig. 5E). Similar to the participant-averaged analysis, individual learning curves show close correspondence between the ideal observer model and the trigram model. Importantly, there are participants where improved predictability of response times does not correspond to adapting a model structure that reflects the real stimulus statistics, but the model underlying response times is still featuring a fully observed structure (Fig. 5G). Note also that the monotonic improvement of CT performance can hinder a richer learning dynamics: several participants have strong non-linearities in their learning as initial improvements correspond to a stronger reliance on a Markov-like structure, which is later abandoned for a more ideal observer-like structure (Fig. 5F and SI Appendix Fig. S9).

We characterized differences between the internal models inferred from individuals on the eighth day of the experiment by swapping the models across subjects and predicting response times using swapped models on the same day. As a control, we performed across-subject swap not only for CT but the Markov model as well. The swapped CT models perform substantially worse than CTs matched to the individuals (two-sided *t*(599) = 25.19, *p* < 0.001, *CI* = [0.0686, 0.0802], *d* = 1.2) and these are outperformed even by the swapped Markov models (two-sided *t*(599) = 5.51, *p* < 0.001, *CI* = [0.00834, 0.0176], *d* = 0.214, Fig. 5J). This latter result indicates that the Markov structure picked up by the CT displays some consistency across participants. We tested whether across-individual differences in the pattern sequences can account for across-individual differences in the internal models by sequence matching (Fig. 5H,I), i.e. regularizing the internal models through inverting the effects of sequence permutations in the iHMM model. The regularized swapped CT performs substantially better than any other swapped model (Fig. 5J) and in particular the regularized Markov model (two-sided *t*(599) = 36.07, *p* < 0.001, *CI* = [0.0702, 0.0783], *d* = 1.26), indicating that individual differences between the models inferred by CT indeed reflect variations in stimulus statistics.

### Trade-off between ideal observer and Markov model contributions

Deviation of the predictive performance of the ideal observer from that of CT highlights that the internal model maintained by participants is only partially accounted for by a model determined by the stimulus statistics. Early-training data also showed that the presence of the ideal observer can be hardly identified. The key question here is whether the margin between CT and ideal observer models can be identified and if it has a structure that is consistent across days within a participant, and whether the structure is consistent across participants. Since initial structure in the internal model reflects inductive biases, this analysis might provide an insight into the nature of inductive biases recruited in a situation when participants are exposed to a new and atypical sequence structure. Since the Markov model has a prominent presence early in the training, we hypothesised that a Markovian structure can make up the difference between CT and the ideal observer model.

First, we tested if the performance of CT can be understood as a combination of the performances of the Markov and ideal observer models. For this, we capitalize on the insight that the Markov and the ideal observer models capture orthogonal aspects of the response statistics. The Markov model can only account for first order transitions across observed stimuli. The ideal observer model is sensitive to both first-order and second-order transitions but since parameters of the ideal observer model are determined by the stimulus statistics, which lack first-order dependencies the structure that this model actually captures is only sensitive to second-order transitions (Table 1). As a consequence, the variances in response times explained by these two models are additive.

We used the additivity of Markov and ideal observer variances to assess how well the performance of CT can be predicted by combining the predictions of the ideal observer and Markov models. We contrasted the normalized CT performance, the difference of the variance explained by CT and the Markov model, with the variance explained by the ideal observer model on a participant by participant basis (Fig. 6A). We found strong correlation between the two measures (*r*(23) = 0.88, *p* < 0.001), indicating that CT performance can be largely explained by a combination of the Markov and ideal observer models. This strong correlation was not only present on the last day of training but during all recording days (*r*(198) = 0.923, *p* < 0.001, SI Appendix Fig. S10).

**Figure 6.**
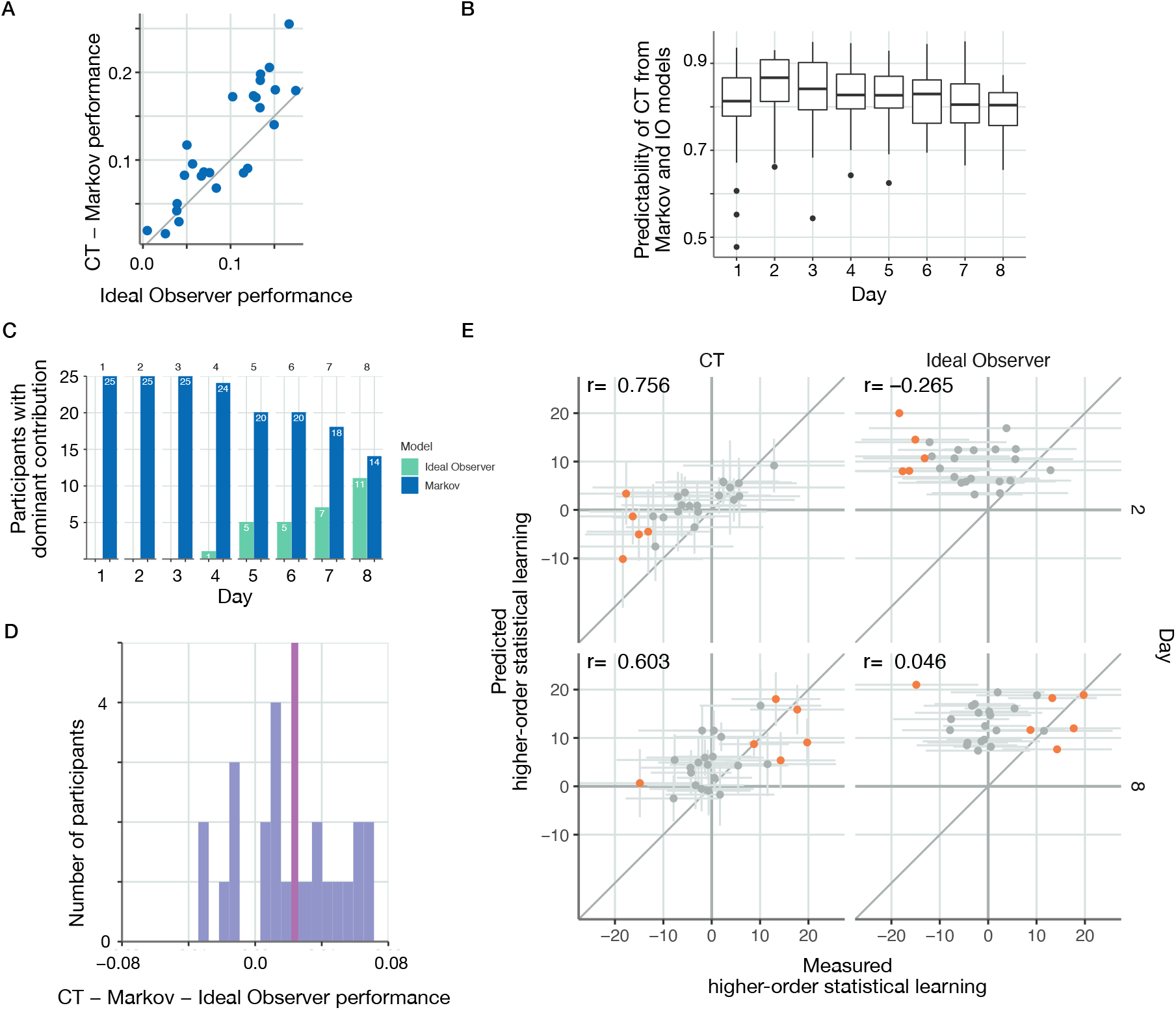
Dissection of CT performance into statistics captured by the ideal observer and Markov models and higher order statistics. **A** Subject-by-subject comparison (*dots* represent individual subjects) of ideal observer model performance and normalized CT performance (the margin by which CT outperforms the Markov model) on Day 8. Dots close to the identity line (*grey line*) indicate cases where CT performance can be reliably accounted for by contributions from the two simpler models. Normalized CT performance closely follows the performance of the ideal observer model, and deviations tend to indicate slightly better normalized CT performance. **B** Performance of a linear model predicting CT model predictions on a trial-by-trial basis from a Markov and ideal observer model predictions on different days of the training. Thick mid-line indicates R^2^ of the trial-by-trial fit of the linear combination to CT performance averaged across participants. Boxes show 25th and 75th percentile of the distribution. **C** Day-by-day comparison of the number of participants for whom the predictive performance of Markov (*blue*) or ideal observer (*yellow*) models was higher. **D** Histogram of the advantage of normalized CT performance over the ideal observer model. Red line marks the mean of the histogram. **E** Higher-order statistical learning in CT (*left panels*) and ideal observer model (*right panels*) on Day 2 (*top panels*) and Day 8 (*bottom panels*) of the experiment. Dots show individual participants. Orange dots represent participants with higher-order learning score significantly deviating from zero. CT can capture both negative deviations (Day 2) and positive deviations (Day 8) in this test and displays significant correlations across participants on both days between the predicted and measured higher-order statistical learning, indicating that subtle and nontrivial statistics of the internal model is represented in CT.

A closer inspection of the response time data can provide exquisite insight into how the inductive bias and evidence-based models are combined to determine responses. In particular, we wanted to assess if trial-by-trial CT predictions can be broken down into the individual contributions of the Markov and ideal observer models. We modelled the response time predicted by CT as a linear combination of the predictions obtained by the Markov and ideal observer models. Response times in all trials of a particular participant for any given day were fitted with three parameters: the weight of the contributing models and an offset. Combined response time predictions showed high level of correlation with the predictions obtained from CT: on any given day the across-participant average correlation was close or above 0.8 (Fig. 6B). Since CT performance can be reliably broken down into the combination of Markov and ideal observer models, a shift in the balance could indicate learning. We assessed the relative contributions of the Markov and ideal observer models by calculating the number of individuals for whom the Markov or the ideal observer model showed higher predictive performance. While the Markov model was more efficient at all but one participant on the first day (Fig. 6C). This prominence shifted towards the ideal observer model by the end of the training (Fig. 6C).

The Markov and ideal observer models capture complementary aspects of stimulus statistics. We investigated if the statistical structure captured by CT goes beyond that captured by the Markov and ideal observer models. The normalised CT showed a small but significant advantage over the ideal observer model on day eight of the experiment (one-sided *t*(24) = 3.646, *p* < 0.001, *CI* = [0.0126, *Inf*], *d* = 0.729, Fig. 6D). Therefore we sought to understand if the marginal advantage of the normalised CT performance reflected relevant stimulus statistics that could be captured by CT but not by the Markov or ideal observer models. We analyzed response times to the third element of three-stimulus sequences which the trigram model is unable to distinguish. In one of the analysed conditions, the first and third elements were pattern elements and we compared these to a condition where the first and third elements are random elements but the actual observations were the same. Since only the latent state differed between the two conditions, these cannot be distinguished by the trigram model. Higher order learning, characterised by response time difference between the two conditions, was highly correlated with the higher-order statistical learning predictions of CT both early in the training (Fig. 6E, *r*(22) = 0.756, *p* < 0.001) and on the last day of training (Fig. 6E, *r*(23) = 0.603, *p* = 0.0014). Interestingly, early in the training most of those participants whose higher-order statistical learning measure was significantly different from zero had negative score (Fig. 6E, orange dots), a counter-intuitive finding termed inverse learning (***Song et al., 2007***; ***Kóbor et al., 2019***). In contrast, higher order statistical learning could not be predicted by the ideal observer (Fig. 6E, *t*(22) = −0.265, *p* = 0.21 and *r*(23) = 0.0457, *p* = 0.828 on Days 2 and 8 of training, respectively).

### Contrasting the internal model inferred by CT to the ideal observer model

Our results show that increased exposure to a complex stimulus statistics results in convergence towards the ideal observer model. Therefore we also wanted to test if there is a closer correspondence between the internal model identified by CT and that represented by the ideal observer. We focused on the patterns in inferences made by the two models, for which we derived a set of relevant measures. First, we used the subjective probability of the stimulus, identical to the measure used to assess response times. Next, entropy of the predictions is used to characterize the uncertainty associated with the stimulus predictions (Fig. 7A). Finally, both models track uncertainty over the latent state of the system continually, which we can measure both before observing a particular stimulus and right after (Fig. 7B). Entropy of state prior characterises the former uncertainty while entropy of state posterior characterises the latter. We calculated these measures for each trial and contrasted them between pattern and random trials both for early models inferred by CT (Day 1) and for models inferred after extensive training (Day 8) (Fig. 7C-F). The ideal observer model describes the way stimuli are generated and therefore has no free parameters. In order to make this model reasonably flexible, we introduced two parameters, which correspond to basic uncertainty in model parameters: the noise level in the dynamics and the noise level in the emission distributions. Changes in these noise parameters merely rescale the measures but leaves the sign of the difference unchanged. The parameters we chose were those that resulted in measures on the same scale as for the CT model. None of the measures derived from the early-training CT models were significantly different from zero. Measures of the late-training CT models showed similar tendencies as the ideal observer model for the subjective probability of the stimulus, state prior and state posterior entropies. It was only the entropy of the predictions which was qualitatively different from the ideal observer model. Taken together, trial-by-trial regressors inferred from response time data provide insights into key quantities of an ideal observer model and have the potential to track these quantities along the training of individuals.

**Figure 7.**
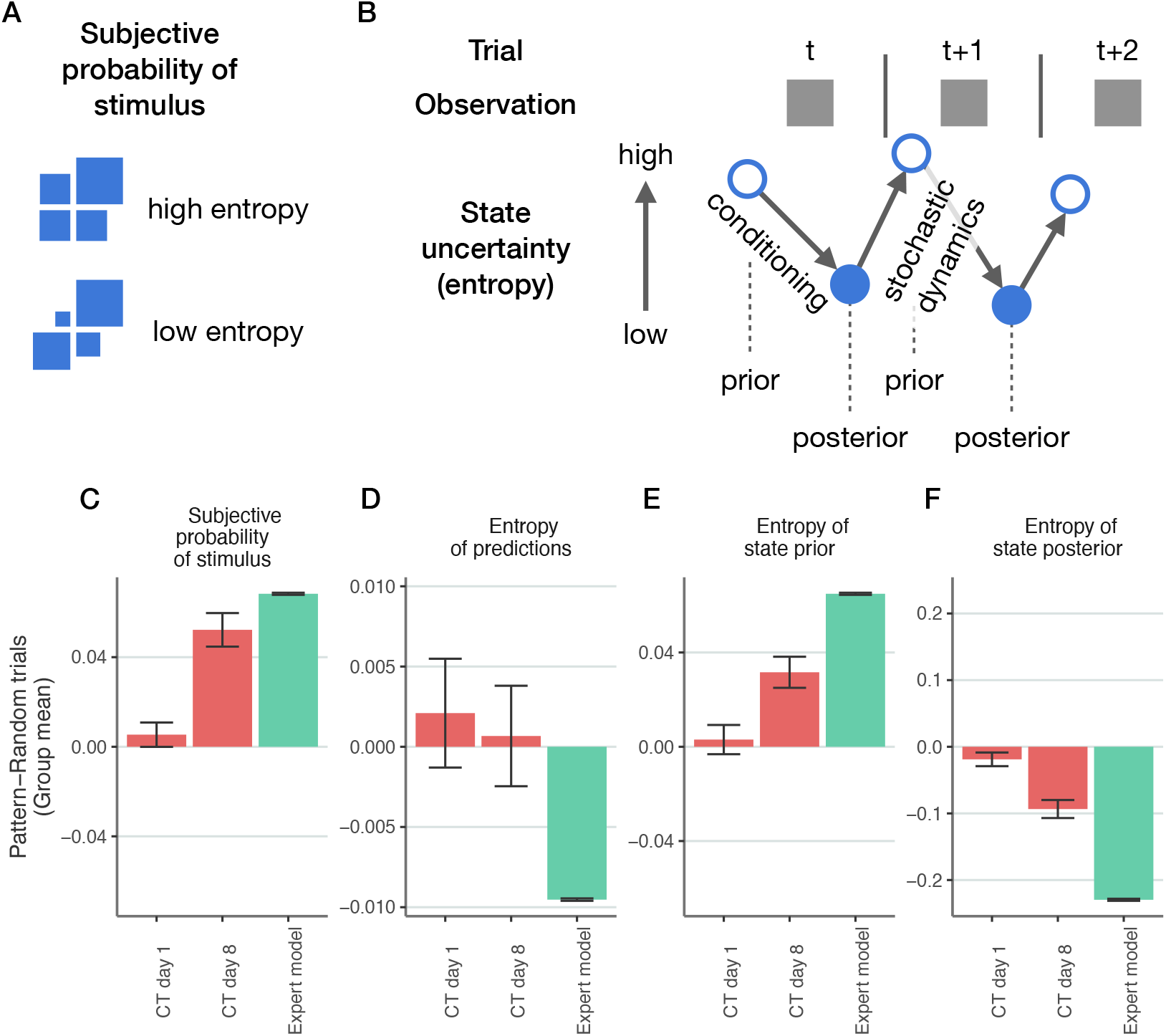
Comparison of the functioning of CT and the ideal observer model. **A**, Illustration of low entropy (*top*) and high entropy (*bottom*) predictive distributions of upcoming stimuli. Size of the square corresponds to the probability of a particular stimulus. **B**, Illustration of latent state entropy dynamics across three consecutive trials. Entropy of the latent state distribution calculated based on previous observations (and thus earlier state estimates) but without the knowledge of the current stimulus (entropy of state prior, *open circles*), and the entropy of the latent state distribution based on both the earlier and the current observations (entropy of the state posterior, *filled circles*). **C-F** We assessed the predictive probability of upcoming stimulus (**C**), the entropy of the predictive probability distribution (**D**), the entropy of state prior (**E**), and entropy of state posterior (**F**) in both CT (*red bars* and the ideal observer model (*green bar*) on individual trials of a stimulus sequence. These four model characteristics were calculated for pattern and random trials separately and the difference between pattern and random trials is shown. CT was evaluated on Day 1 and Day 8 of the training as well. For the ideal observer model, it is the deviation from zero and direction of deviation that are the relevant quantities since the scale is determined by two noise parameters of the model and is therefore arbitrary. Bars denote mean differences between pattern and random trial means across trials and across models inferred for different participants, and error bars show 2 s.e.m.

## Discussion

In this paper we have built on the idea of Cognitive Tomography, which aims to reverse engineer internal models by separating the task-general component from the task-specific component of a generative model of behaviour, to infer high-dimensional dynamical internal models. Key to our approach was the combination of non-parametric Bayesian methods that allow discovering flexible latent variable models, with probabilistic programming, which allows efficient inference in probabilistic models. The proposed approach yields a tool, which has a number of appealing properties for studying how acquired knowledge about a specific domain affects momentary decisions of biological agents: 1, We used iHMM, a dynamical probabilistic model that can naturally accommodate rich inter-trial dependencies, a characteristic prior of humans (***Glaze et al., 2015***; ***Urai et al., 2017***; ***Talluri et al., 2018***); 2, iHMM has the additional benefit of being able to capture arbitrarily complex statistical structure but not increasing the complexity of the model more than necessary (***Gershman and Blei, 2012***; ***MacKay, 2003***); 3, We could predict response times of human participants on a trial-by-trial basis; 4, Complex individualized internal models could be inferred from behavioural data; 5, We evaluated individual learning curves, and thus could contrast the learned internal model to the ideal observer model that describes the ground truth of how stimuli were generated. Cognitive Tomography revealed considerable deviations from the ideal observer model and instead transient models were identified at individuals that were nurtured temporarily only to be abandoned later during training. We used CT to investigate the inductive biases of participants. We demonstrated that the internal model discovered by CT can be reliably broken down into two components: 1, a Markovian component, which only captures immediate dependence on the preceding observation; 2, and the ideal observer model. The latter component can be identified with the evidence participants collected about the true stimulus statistics and the former component can be identified with an inductive bias participants relied on early in the training. Indeed, while there were individual differences in how long the Markovian hypothesis about stimulus structure was maintained, the evidence-based component grew in dominance with more extensive training. Importantly, we have validated that the statistical structure posited by Cognitive Tomography reflects key characteristics of internal models: specificity to environmental statistics and generality over tasks. For this, we dissociated the contributions of the two main components of the learned model: the behavioral model was replaced for predicting erroneous responses of participants, and adaptation to stimulus statistics was demonstrated by differential recruitment of the internal model under different stimulus statistics.

Learning in general is an ill-defined, under-determined problem. Learning requires inductive biases formulated as priors in Bayesian models to efficiently support the acquisition of models underlying the data (***Griffiths et al., 2010b***; ***Mitchell, 1980***). The nature of such inductive biases is a fundamental question which concerns both cognitive science and neuroscience, even machine learning (***Richards et al., 2019***; ***Griffiths et al., 2010b***; ***Botvinick et al., 2019***). These inductive biases determine what we can learn. Variance in inductive biases across individuals leads to different learning strategies, and can be tightly linked to the neural representations of the models. The CT approach helps identifying individualized internal models from behavioral data thus allowing the assessment of individual learning strategies. Key to this approach is data efficiency: the experimenter needs to infer internal models from limited data. Bayesian inference that exploits structured models offers exactly this data efficiency while maintaining flexibility to accommodate potentially complex, high-dimensional models (***Ghahramani, 2015***). Our analysis identified two components that dominated the internal model learned by CT: the Markov model and the ideal observer model. Since the novel statistics governing the stimuli is captured by the ideal observer model, the Markov model could be identified with an inductive bias that helps learning when data is scarce, i.e. early in the training. Inductive biases can effectively support learning if these represent priors, which reflect the statistics of the environment. Indeed, Markovian dynamics can be a good approximation of the dynamics of the natural environment therefore can constitute a useful inductive bias. Dependence of response time on the immediately preceding response time could emerge purely from physiological constraints, such as the interdependence of finger movements. We believe that the learning curves revealed by CT support that the Markovian structure is indeed a result of the internal model rather than such constraints because the Markovian component of some participants completely diminishes after a number of days of training. A minor contribution of these factors, however, cannot be excluded solely based on response time data, but our analysis provides a strong support that initial inductive bias is dominated by a Markov model. Taken together, the prevalence of a Markovian component in the internal model indicates that humans rely on a simple dynamical inductive bias when faced with a novel sequential statistics.

The presented model builds on the original CT analysis performed on faces (***Houlsby et al., 2013***) but differs in a number of fundamental ways. We sought to infer the evolution of the internal model for statistics new to participants. In contrast to the earlier formulation using a 2-dimensional latent space and a static model, here the inference of a dynamical and potentially high-dimensional model yields a much richer insight into the working of the internal model acquired by humans. Using a structured internal model allows the direct testing of the model against alternatives, thus providing opportunities to reveal the computational constraints that might limit learning and inference in individuals. A well-structured internal model can be used to make arbitrary domain-related inferences within the same model. Based on this, we can decompose the complex inference problem into separately meaningful sub-parts which can be reused in tangential inference problems to serve multiple goals. By showing that the same variables can be used for multiple tasks, it is reasonable to look for signatures of these quantities in neural representations. A possible alternative formalization of this problem could be using Partially Observed Markov Decision Process (***Wu et al., 2018***), where internal reward structure and the subjective belief of the effect of the participant’s actions are jointly inferred with the internal model. However, in our experiment, the action model has a simple structure and hence the problem simplifies to a probabilistic sequence learning problem. Instead, here we focus on inferring rich internal model structures as well as having an approximate Bayesian estimate instead of point estimates as in ***Wu et al. 2018***.

Our model produces moment by moment regressors for (potentially unobserved) variables that are cognitively relevant. Earlier work considered neural correlates of hidden state representations in the orbitofrontal cortex of humans (***Schuck et al., 2016***) but the internal model was not inferred, rather assumed to be fully known. CT provides an opportunity to design regressors for individualised and potentially changing internal models. In particular, the model differentiates between objective and subjective uncertainties, characteristics relevant to relate cognitive variables to neural responses (***Barthelmé and Mamassian, 2009***; ***Bach and Dolan, 2012***; ***Michael et al., 2015***; ***Pouget et al., 2016***). The former is akin to a dice-throw, uncertainty about future outcomes which may not be reduced with more information. The latter is uncertainty arising from ambiguity and lack of information about the true current state of the environment. We showed that uncertainties exhibited by a trained individual’s internal model show similar patterns in these characteristics as the ideal observer model, which promises that uncertainties inferred at intermediate stages of learning are meaningful.

Recently, major efforts have been devoted to learning structured models of complex data both in machine learning and in cognitive science (***Lake et al., 2015***; ***Kemp and Tenenbaum, 2008***; ***Battaglia et al., 2013***; ***Saxe et al., 2019***). These problems are as diverse as learning to learn (***Lake et al., 2015***; ***Braun et al., 2010***), causal learning (***Goodman et al., 2011***), learning flexible representational structures (***Kemp and Tenenbaum, 2008***), visual learning (***Austerweil and Griffiths, 2013***). When applied to human data to reverse engineer the internal models harnessed by humans, past efforts fall into two major categories. 1, Complex (multidimensional) models are inferred from data and fitted to across-participant averaged data (***Orbán et al., 2008***; ***Griffiths and Tenenbaum, 2006***; ***Battaglia et al., 2013***), ignoring individual differences. 2, Simple (low dimensional) models are used to predict performance on a participant-by-participant manner, thus resulting in subjective internal models (***Gold and Stocker, 2017***; ***Glaze et al., 2015***). In particular in a simple two-latent variable dynamical probabilistic model individualised priors have been identified (***Glaze et al., 2018***). In this setting binary decisions were sufficient as an ‘expert model’ was assumed and assessment of prior comprised of inferring a single parameter, which defined the width of a one-dimensional hypothesis space. Findings of this study gave insights into how individuals differ in their capacity to adapt to new situations. Recently, a notable approach has been presented, which aims at characterizing individual strategies in a setting where the complexity of the state space is relatively large (***van Opheusden et al., 2021***). In this study, the rules of the game (equivalent of the statistics of stimuli in our case) and the relevant features (equivalent to the latent variables in our case) were assumed to be known by the participants. However, being a two-player task there was uncertainty about the strategy of the opponent and the limitations in the computational complexity of the inference was investigated. This aspect is orthogonal to the aspects investigated here and therefore highlight additional appeal of analysing behavior in complex settings. The contribution of the current paper is twofold: 1, We exploit recent advances in machine learning to solve the reverse-engineering problem in a setting where complex internal models with high-dimensional latent spaces are required; 2, We contribute to the problem of identifying structured inductive biases by enabling direct access to the internal model learned by individuals and by dissecting the contributions of evidence and inductive bias.

A widely studied approach to link response time to quantities relevant to task execution is the drift diffusion model, DDM (***Ratcliff et al., 2016***). In its most basic form evidence is stochastically accumulated as time passes such that the rate of accumulation is proportional to the information gained by extended exposure to a stimuli, until evidence reaches a bound where decision is made. Through a compact set of parameters DDM can explain a range of behavioural phenomena, such as decisions under variations in perceptual variables, adaptation to the volatility of the environment, attentional effects on decision making, the contribution of memory processes to decision making, decision making under time pressure (***Drugowitsch et al., 2012***; ***Nosofsky et al., 2011***; ***Palmer et al., 2005***; ***Smith and Ratcliff, 2009***; ***Ossmy et al., 2013***), and neuronal activity was also shown to display strong correlation with model variables (***Gold and Shadlen, 2001***; ***Hanes and Schall, 1996***). Both LATER and DDM have the potential to incorporate variables relevant to make decisions under uncertainty and the marginal distributions predicted by the two models are comparable. Our choice to use the LATER model was motivated by two major factors. First, LATER is formulated with explicit representation of subjective predictive probability by mapping it onto a single variable of the model. This setting promises that subjective probability can be independently inferred from available data and the internal model influences a single parameter of the model. As a consequence, subjective probability is formally disentangled from other parameters affecting response times and associated uncertainty can be captured with Bayesian inference. In case of distributing the effect of subjective probability among more than one parameters (starting point, slope, variance) the joint inference of subjective probability with other parameters affecting response times results in correlated distributions. Consequently, maximum likelihood inference, or any other point estimations, the preferred method to fit DDM, will have large uncertainty over the true parameters due to interactions between other variables. Furthermore, this uncertainty remains unnoticed as there is usually no estimation of this uncertainty, only point estimates. Second, trials are usually sorted based on the design of the experiment into more and less predictable trials (with notable exceptions like (***Glaze et al., 2015***)). This leads to a misalignment between the true subjective probabilities of a naive participant and the experimenter’s assumptions. Assuming full knowledge of the task and therefore assuming an impeccable internal model in more complex tasks, however, implies that potential variance in the acquired internal models across subjects will be captured in variances in parameters characteristic of the response time model rather than those of the internal model. DDM is considered to be an algorithmic-level model (***Marr, 1982***) of choices (***Talluri et al., 2018***), which is indeed useful for linking choice behaviour to neuronal responses (***Gold and Shadlen, 2007***). The appeal of the Bayesian description offered by the normative framework used here is that it can accommodate a flexible class of internal models, without the need to adopt algorithmic constraints. Similar algorithmic-level models of behaviour that is based on the flexible and complex internal models yielded by Cognitive Tomography are not available and will be the subject of future research.

## Acknowledgements

This research was supported by the National Brain Research Program (project 2017-1.2.1-NKP-2017-00002, D.N., G.O.); Hungarian Scientific Research Fund (NKFIH-OTKA K K125343, G.O.; NKFIH-OTKA K 128016, D.N., NKFIH-OTKA PD 124148, K.J.);Janos Bolyai Research Fellowship of the Hungarian Academy of Sciences (K.J.); IDEXLYON Fellowship of the University of Lyon as part of the Programme Investissements d’Avenir (ANR-16-IDEX-0005) (D.N). B.T. was supported by scholarship by Budapest University of Technology and Economics as well as by Mozaik Education Ltd. (Szeged, Hungary). Resources for the computational analysis were generously provided by the Wigner Data Center. The authors would like to thank to Peter Dayan and Noémi Éltető for comments on an earlier version of the manuscript.

## Author contributions

B.T., D.G.N. and G.O. designed the computational model. M.K., K.J., D.N. designed the experiment. M.K. recorded the experimental sessions. B.T. implemented the models and carried out analyses. B.T., G.O., D.G.N., K.J., D.N. interpreted results. B.T. and G.O. wrote the manuscript and the supplementary information. D.G.N., K.J., D.N. commented on the manuscript at all stages.

## Experiment: Materialsand methods

### Participants

Twenty-five individuals (22 females and 3 males) aged between 18 and 22 (*M_Age_* = 20.4 years, *SD_Age_* = 1.0 years) took part in the experiment (we recruited 32 participants, but only 26 completed the experiment; we omitted one further participant because of a system error which resulted in partial loss of their experiment data). They were university students (*M_Years of education_* = 13.3 years, *SD_education_* = 1.0 years) from Budapest, Hungary. None of the participants reported history of developmental, psychiatric, neurological or sleep disorders, and they had normal or corrected-to-normal vision. They performed in the normal range on standard neuropsychological tests of short-term and working memory (Digit span task: *M* = 6.48, *SD* = 1.15, Counting span task: *M* = 3.76, *SD* = 0.99) (***Janacsek and Nemeth, 2013***). Before the assessment, all participants gave signed informed consent and received course credit for participation. The study was approved by the Institutional Review Board of Eötvös Loránd University, Hungary.

### Tasks

Alternating Serial Reaction Time (ASRT) Task Learning was measured by the ASRT task (***Howard and Howard, 1997***; ***Nemeth et al., 2010***). In this task, a stimulus (a dog’s head) appeared in one of four horizontally arranged empty circles on the screen and participants were asked to press the corresponding button as quickly and accurately as they could when the stimulus occurred. The computer was equipped with a keyboard with four heightened keys (Z, C, B, M on a QWERTY keyboard), each corresponding to a circle in a horizontal arrangement. Participants were asked to respond to the stimuli using their middle- and index fingers bimanually. The stimulus remained on the screen until the participant pressed the correct button. The next stimulus appeared after a 120 ms response-to-stimulus-interval (RSI). The task was presented in blocks of 85 stimuli: unbeknownst to the participants, after the first five warm-up trials consisting of random stimuli, an 8-element alternating sequence was presented ten times (e.g., 2r4r3r1r, where each number represents one of the four circles on the screen and r represents a randomly selected circle out of the four possible ones).

### Procedure

There were ten sessions in the experiment, with one-week delay between the consecutive sessions. Participants performed the ASRT task with the same sequence in the first eight sessions, then an interfering sequence was introduced in Session 9, and both (original and interfering) sequences were tested in Session 10 (see S1). Participants were not given any information about the regularity that was embedded in the task in any of the sessions (***Nemeth et al., 2010***). They were informed that the main aim of the study was to test how extended practice affected performance on a simple reaction time task. Therefore, we emphasized performing the task as accurately and as fast as they could. Between blocks, the participants received feedback about their average accuracy and reaction time presented on the screen, and then they had a rest period of between 10 and 20 s before starting the next block. On Days 1-9, the ASRT consisted of 25 blocks. One block took about 1-1.5 min, therefore the task took approximately 30 min. For each participant, one of the six unique permutations of the four possible ASRT sequence stimuli was selected in a pseudo-random manner (***Howard and Howard, 1997***; ***Kóbor et al., 2017***; ***Nemeth et al., 2010***). The ASRT task was performed with the same pattern sequence in Sessions 1-8. In Session 9, the ASRT was performed with a new interfering pattern sequence. In Session 10, participants performed 20 blocks of the ASRT task switching between the pattern sequences of Sessions 1-8 and Session 9 every five blocks. In Session 10, the task took approximately 24 min. After performing the ASRT task in Session 10, we tested the amount of explicit knowledge the participants acquired about the task with a short questionnaire. This short questionnaire (***Nemeth et al., 2010***; ***Song et al., 2007***) included two questions: “Have you noticed anything special regarding the task?” and “Have you noticed some regularity in the sequence of stimuli?”. The participants did not discover the true probabilistic sequence structure.

**Figure S1.**
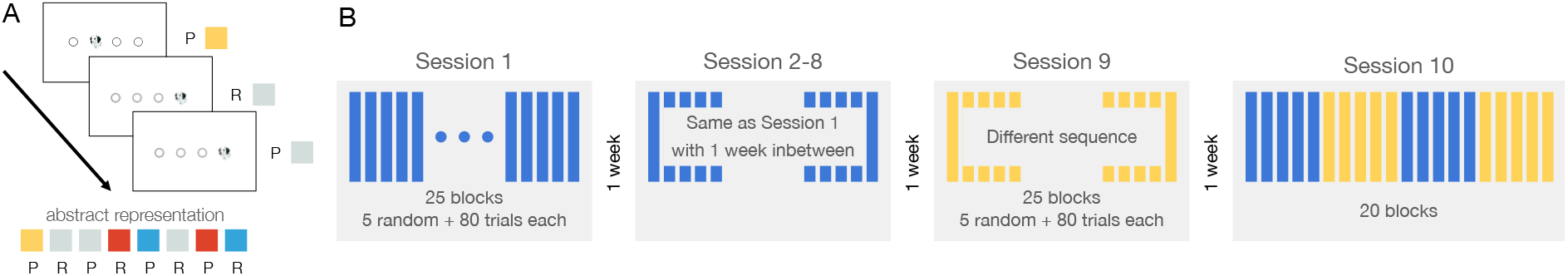
*A* Experimental stimuli and abstract representation used in the paper. *B* Design of the experiment. The experiment consisted often sessions, separated by a one-week delay. On Days 1-8, participants performed the ASRT task with sequence 1 throughout 25 blocks (5 epochs) each sessions. On Day 9, an interfering sequence (sequence 2) was introduced. Both sequences were tested on Day 10 with blocks of 5 alternating.

## Background

### Models for Sequential Prediction

The experimental stimuli form a sequence of discrete observations in discrete time, 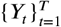. The task is therefore to predict the upcoming stimulus conditioned on the history of observations:

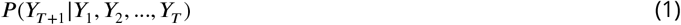

In practical terms, learning a model for this temporal prediction task requires imposing a structure over these conditional distributions. Without structural assumptions, there is no statistical dependence among different histories, that is, there is no generalisation from history to future observations.

In the following section we introduce a computational model, the Hidden Markov Model, which can provide a general language for solutions of this problem. It can express arbitrarily complex models given sufficiently large amounts of data. In order to remain as general as possible, we will consider a model space (infinite Hidden Markov Models as in ***Gael et al. 2008***) which can model all the possible distributions in equation 1. Moreover, we would like to achieve this while being able to express inductive biases in this language which are useful for constraining the possible models in the limited data case.

### Hidden Markov Model

Formally, a Hidden Markov Model comprises of a sequence of hidden states 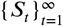 and a sequence of observations 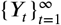. In this work we take both the latent states and the observations to be discrete, that is *S_i_*, *Y_i_*, ∈ ℕ sequence of hidden (latent) states constitute a discrete Markov-chain with transition probabilities *π_ij_* = *P*(*S*_*t*+1_ = *j*|*S_i_* = *i*). In a Markov-chain, the sequence element *S_t_* is conditionally independent of the history conditioned on the previous state and the transition probabilities:

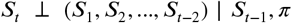

At (discrete) time *t*, observation *Y_t_* is governed by the latent state *S_t_*. The observations are generated independently and identically, conditioned on the (latent) state:

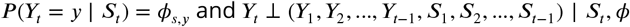

Importantly, since the latent state can incorporate arbitrary information (identical observations at different time-points can correspond to different states), assuming arbitrarily many latent states, we get a completely general solution for the prediction problem in 1. With an adequate prior (e.g. the Hierarchical Dirichlet Process in ***Teh et al. 2006*** we can learn such structures efficiently (***Gael et al., 2008***)). In practical terms this means the assumed number (e.g. posterior mean number) of different states increases with the length of the observation sequence. Loosely: until proven otherwise, a simpler structure is assumed.

## Cognitive Tomography

We construct a model of behaviour which consists of two parts:

1. An internal model maintained by the participant, which formalizes how latent states assumed to underlie observations evolve and how these states are linked to observations.
2. A model relating the prediction of participants’ internal model to their responses (response time model).

### Doubly Bayesian Model

Due to the uncertainty of the participants about the true model and actual state of the stimuli and to the uncertainty of the experimenter about the model maintained by participants and about the actual state of this internal model, the problem can be described as doubly Bayesian. We do Bayesian inference over an internal representation of individuals who themselves do Bayesian inference. Elements of the experimenter’s model are introduced in following sections.

Prediction of response times can be described by the following algorithm:

1. We take posterior samples from the behavioural model which consists of parameters of the internal model and the response time model conditioned on data from ten consecutive blocks of trials, where:

a. all stimuli, and
b. response times (with incorrect trials’ first five random trials’ response times, and response times smaller than 180 msec in each block removed). According to the original formulation by ***Carpenter and Williams 1995***, fast response times come from an alternative distribution. We cut off the fast response times (as in ***Kim et al. 2017***) at the fixed 180 msec value. However, we did not fit the cut-off time parameter.
are included.
2. For each of the 60 posterior model samples we compute predicted response times by:
  a. filtering the belief over the latent state over the entire sequence
  b. produce subjective probabilities for each trial
  c. produce response time prediction (MAP estimate conditioned on the subjective probability and the response time parameters of the model sample)
Then we marginalize (i.e. average) over the response time predictions of model samples.
3. We evaluate model performance by computing the *R*^2^ explained variance measure of the predicted response times on the response times of the test dataset.

Note: since actual beliefs depend on past beliefs, one can think of the belief sequence as the path of a light-ray in a large dimensional fog (representing the state uncertainty). During inference, we have a noisy measurement of the light-ray in different points of time and we would like to reconstruct the best explanation of the observation sequence (response times) in terms of a hidden path. As for prediction, the model produces response time predictions for the entire stimulus sequence with no further feedback of response times (i.e. estimated internal beliefs are not updated based on what response time the participant produced on given trials).

### Infinite Hidden Markov Model

The infinite Hidden Markov Model is a non-parametric extension of the Hidden Markov Model, assuming infinitely many states. There is a hierarchical prior imposed over the state transition matrix and the so-called emission distributions relating the latent (hidden) states to observations (Fig. S2).

The hierarchical prior we used is exactly the one defined in ***Gael et al. 2008***. We extended their implementation of their model to a doubly Bayesian behavioural model including the response time.

#### Internal model of participants

A participant is assumed to learn a probabilistic model of the sequence which is formalized as an infinite Hidden Markov Model. At (discrete) time *t*, observation *Y_t_* is governed by a latent (not directly observable) state *S_t_*. The states {*S_t_*}_*t*=1,2,…_ constitute a Markov-chain, which means the following:

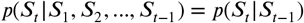

That is, the state *S*_*t*−1_ holds all information about past regarding the possible evolution of system. In other terms, conditioning on state *S*_*t*–1_ renders *S_t_* and all previous states *S*_1_,*S*_2_,…, *S*_*t*–2_ statistically independent.

The observation *Y_t_* at time *t* is independent of all other observations, conditioned on the latent state *S_t_* (and the model parameters). That is, once the state of the system is decided, the actual previous observations are independent of *Y_t_*.

The parameters governing the state transitions are aggregated in the parameter matrix *π*:

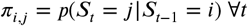

The observation distributions are given by the parameter matrix *ϕ*:

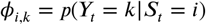

At any given time during the task, we assume the participant had estimated the parameters *π* and *ϕ* and uses these (point estimates) to do exact filtering over the sequence of observations. That is, in each trial they use the evidence provided by the current stimulus to update their belief over the latent state of the sequence. When doing computations with the participant’s internal model, we hold the internal model fixed within shorter time-scales of the task (e.g. one session). The participant represents their belief about the current latent state of the system by a posterior distribution, updated by each incoming observation, while always conditioning on their current estimates 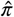 and 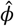 of *π* and *ϕ* respectively.

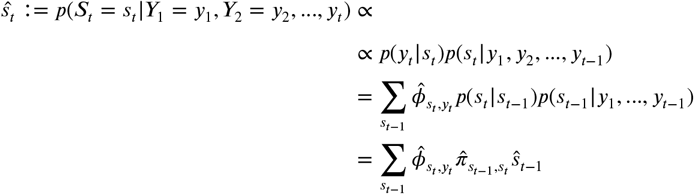

For filtering, stimuli of all trials (including initial random trials at the beginning of each block and stimuli in trials where participant hit the wrong key initially) are used. That is, even if response times are not considered when doing inference over the participant’s internal model, the participant is assumed to update their internal beliefs based on the stimulus shown.

Prediction of the next stimulus is computed by marginalizing over the latent state posterior distribution:

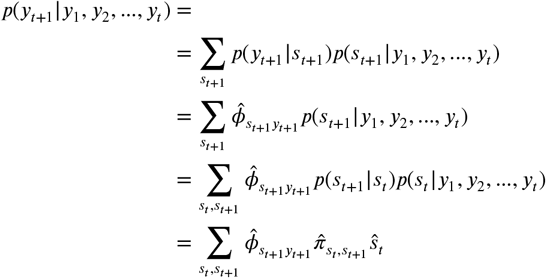

#### Participant learning the model

Throughout the execution of the task, the internal model of the participants is continually updating. We do not directly model the computation of the participants that estimates the current *π* and *ϕ* parameters. That is, within a given train or test dataset (10 consecutive blocks) we hold *π* and *ϕ* fixed. We do allow, however, for these estimates of *π* and *ϕ* to change between sessions.

#### Inference

We do Approximate Bayesian Inference using a custom sampling method that mixes steps of a Hamiltonian Monte Carlo (HMC) and a Gibbs sampler which samples a slicing parameter (see ***Gael et al. 2008***). The priors used in the model are listed in Table 2.

In order to handle the infinitely many possible states, we use a modified version of the slice sampling method described in (***Gael et al., 2008***). In the original beam sampling algorithm, the authors sample the latent state sequence and make use of the slicing variable to constrain the set of used states to a finite set. They sample the latent sequence and the slicing variables in an alternating fashion. In our case we do not sample latent state sequences, instead, we have to estimate the subjective belief sequence over the latent states. In this latter case, the posterior belief is infinite dimensional and we use slicing to approximate this infinite-dimensional computation with a finite one. At each sampling step, we only look at the latent state belief distribution’s 1 – *ϵ* support where *ε* is sampled from Uniform(0.02, 0.2).

**Figure S2.**
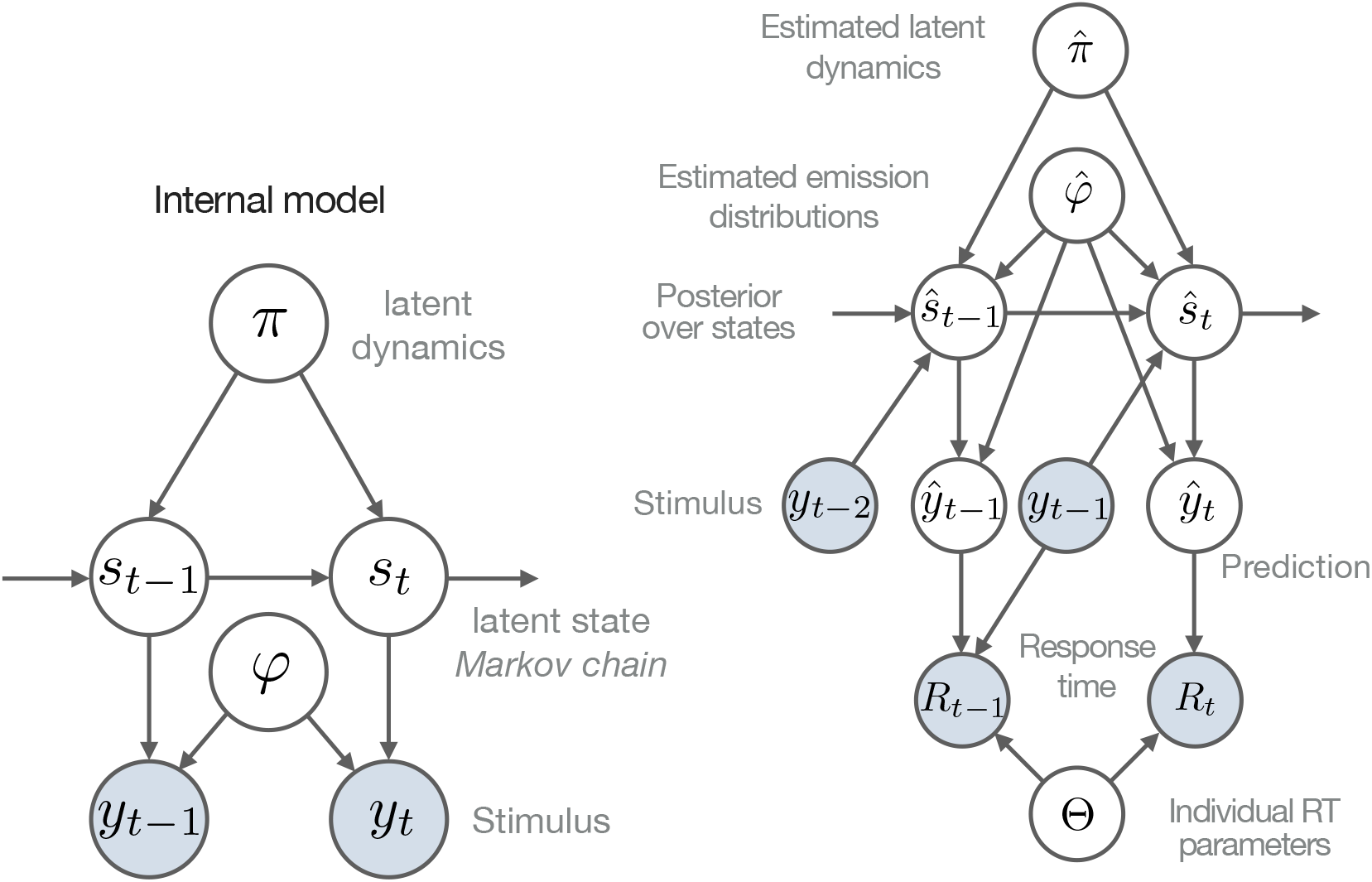
Graphical representation of internal model and generative model of behaviour. Left: Internal model, generative model of the sequence assumed by the participant. Right: generative model of behaviour.

**Table 1.** Summary of when model parameters are allowed to change.

| Variable | Notation | Within train/test | Within session, between train-test | Between sessions | Between participants |
| --- | --- | --- | --- | --- | --- |
| State transition distribution | $\hat{\pi}$ | No | No | Yes | Yes |
| Observation distribution | $\hat{\varphi}$ | No | No | Yes | Yes |
| State belief | $\hat{s}_t$ | Yes | Yes | Yes | Yes |
| Response time parameters | $\tau_0, \mu, \sigma$ | No | No | No | No |
| Prior of observation distribution | $H$ | No | No | No | No |
| Hierarchical prior over state transitions | $\alpha, \beta$ | No | No | No | No |

**Table 2.** Parameter priors. Values of the hierarchical prior over state transitions taken from [(***Gael et al., 2008***)]

| Variable | Prior |
| --- | --- |
| State transition distribution | $\hat{\pi}_i \sim \text{Dirichlet}(\alpha_0/K, \dots, \alpha_0/K, \alpha_0/K \cdot \epsilon)$ |
| Observation distribution | $\hat{\phi} \sim \text{Dirichlet}(0.8, 0.8, 0.8, 0.8)$ |
| Response time parameters | $\tau_0 \sim \Gamma(1, 10)$<br>$\mu \sim \Gamma(1, 0.1)$<br>$\sigma \sim \Gamma(1, 0.01)$ |
| Hierarchical prior over state transitions | $\alpha = 1.3$<br>$\gamma = 3.8$ |

Four independently and randomly initialised Markov Chains were sampled with 1600 steps of the slice sampling (outer Gibbs-sampling chain) and 30 NUTS steps were taken inbetween the slice sampling steps each time. Samples from the second half of each chain were used to check if estimates of response time parameter means and confidence intervals were identical. For prediction, last 60 unique samples were used from each chain because prediction performance saturates at this number of samples.

### Ideal Observer Model

#### Internal Model Components

We formalise the ideal observer the following way: at any given point of the experiment, the ideal observer entertains an internal dynamical model comprising of two parts: latent dynamics (the transition probabilities between latent states) and an observational model (conditional distributions of observations conditioned on the latent state).

##### Filtering

In order to produce predictions for the upcoming observation, conditioning on a fixed model, the ideal observer solves the filtering problem:

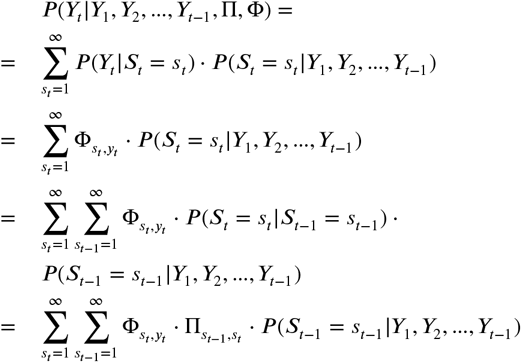

The term filtering is used because as we deduced, the relevant quantity is *P*(*S*_*t*-1_ |*Y*_1_,*Y*_2_,…,*Y*_*t*-1_) which can be filtered through our observations. We carry on this quantity and can calculate it for the next time-step using our model parameters and the observation *Y_t_*.

Importantly, instead of sampling one possible latent trajectory, we have to marginalise over these latent sequences to obtain our prediction for the upcoming stimulus. That is, our prediction is the aggregate of the predictions many possible latent pasts. We combine the predictions of ‘had these been the sequence of causes of my past experiences, I should see this’ for all possible hypothesised latent cause sequences.

#### Response Time Model

In order to connect the predictions of the internal model to measured behaviour, we need to employ a generative model of response times in the form of a conditional probability distribution conditioned on the subjective predicted probability of the upcoming stimulus. To achieve this, we employ the reaction time model of ***Carpenter and Williams 1995***, which in its original formulation states that the majority of saccadic response times come from a reciprocal Normal distribution.

Further studies suggest choice response time distribution should have a similar form (***Brown and Heathcote, 2008***; ***Harris et al., 2014***). However, in other formulations, there is no explicit dependence of the distribution of the RT in a single trial depending on the subjective predicted probability, hence those models are inadequate for our purposes. The generative model for correct response times (LATER model, ***Carpenter and Williams 1995***) is:

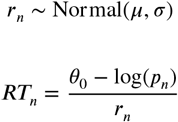

where *p_n_* is the subjective probability (output of the internal model) corresponding to the actual upcoming stimulus and *μ,σ,θ*_0_ are the parameters characterising an individual’s response time model. These parameters jointly describe the mean and variance of the response times. Note that in our experiment these parameters comprise all idiosyncratic effects at hand, namely the individual’s state, their response times’ sensitivity to subjective predicted probabilities, the effects of instruction influencing speed-accuracy trade-off.

The response time parameters are jointly inferred along with the internal representations (dynamical model, observation distribution, latent state inference).

#### Validation on synthetic datasets

In order to validate our behavioural model as well as our inference method, we looked at how well we can recover subjective probabilities on a synthetic dataset. We used the algorithm in ***Gael et al. 2008*** on synthetic ASRT data to infer a first set of three different internal models from different levels of exposure. These models represent internal models of different synthetic participants (Fig. S3A). We take these models as the ground truth for our synthetic experiment. We trained one model on 640, 1280 and 2400 trials of ASRT stimuli. We then generated response times from the generative model with three parameter settings for each of *τ*_0_, *μ*, and *σ* resulting in a total of 3^3^ = 27 different synthetic response time sequences. The resulting response time distribution’s variance is influenced by all four factors - the subjective probabilities (which depends on the internal model) and the three response time parameters. The standard deviation of the response times is an appropriate measure since this can be also computed for data obtained from human participants. We generated the response times for 10 ASRT blocks (the same number we used for inference on human data). Standard deviation of the resulting response times (*symbol colours* on SI Appendix Fig. S3B) arise from the interaction of all parameters. Different combinations of the response time parameters resulting in the same standard deviation are marked by identical symbols. Then, we used the CT inference method to generate a second set of (posterior) internal model samples. We computed the same model performance measure as for human data (response time prediction performance) and also the performances for subjective probability predictions (SI Appendix Fig. S3B). The results show that the prediction performance of the subjective probabilities exceeds that of the individual response times. Also, as seen on SI Appendix Fig. S3C, standard deviation of human participant’s response times are considerably below the threshold where our CT inference method in the synthetic data experiment was successful.

## Alternative models

### Markov model

#### Internal model of participants

According to this model, the participants assume that the sequence of observations constitute a Markov-chain. That is, for the sequence of observations *y_t_*, we have

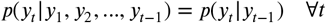

The above equation states that the next observation is independent of all previous observations given the previous observation. This is equivalent to saying that all information (besides parameters governing the sequence) about the state of the sequence is included in the previous observation.

#### Inference

We use the same parameter priors for the response time model as for the iHMM model and the prior for transition probabilities *π_i_* ~ Dirichlet(*α*_0_/*K*, *α*_0_/*K*, *α*_0_/*K*, *α*_0_/*K*), where *K* is the number of states, in this case 4.

Four independently and randomly initialised Markov Chains were sampled with 1600 steps taken with the NUTS sampler in STAN. Samples from the second half of each chain were used to check if response time parameter estimates’ means and confidence intervals were identical. For prediction, last 60 samples were used from each chain.

#### Relation to sequential motor effects

The prediction a Markov internal model makes can be rephrased as “how quick a participant will respond to the next stimulus *j* if the current stimulus is *i*”. In our paradigm, the mapping between the stimuli and the required responses are fixed. For this reason, predictions of a Markov model are indistinguishable from predictions made by a “sequential motor bias” model positing a dependency of the response time on which successive key strokes are made (i.e. which finger the participant uses for the current trial and which one for the previous trial).

Our investigation implies that the Markov models inferred are a superposition of sequential motor effects and an internal model that is Markovian (see Results for our argument).

#### Relation to HMM

Note that Markov models are subset of Hidden Markov Models. We can always write a Markov model as an HMM if we have a matching number of observation values and latent state values and each observation is unique to a state.

This is particularly important since for an HMM for which the above condition holds, there is an equivalent Markov chain that describes the exact same sequence structure. This is the reason why we term some of the internal models identified by our iHMM method “Markov-like”, since they are closely approximated by an actual Markov model.

### Trigram model

The model we describe here is also referred to as ‘triplet model’ in previous works using the ASRT paradigm. We use the term trigram since it is more commonly used in a sequential prediction modelling context.

**Figure S3.**
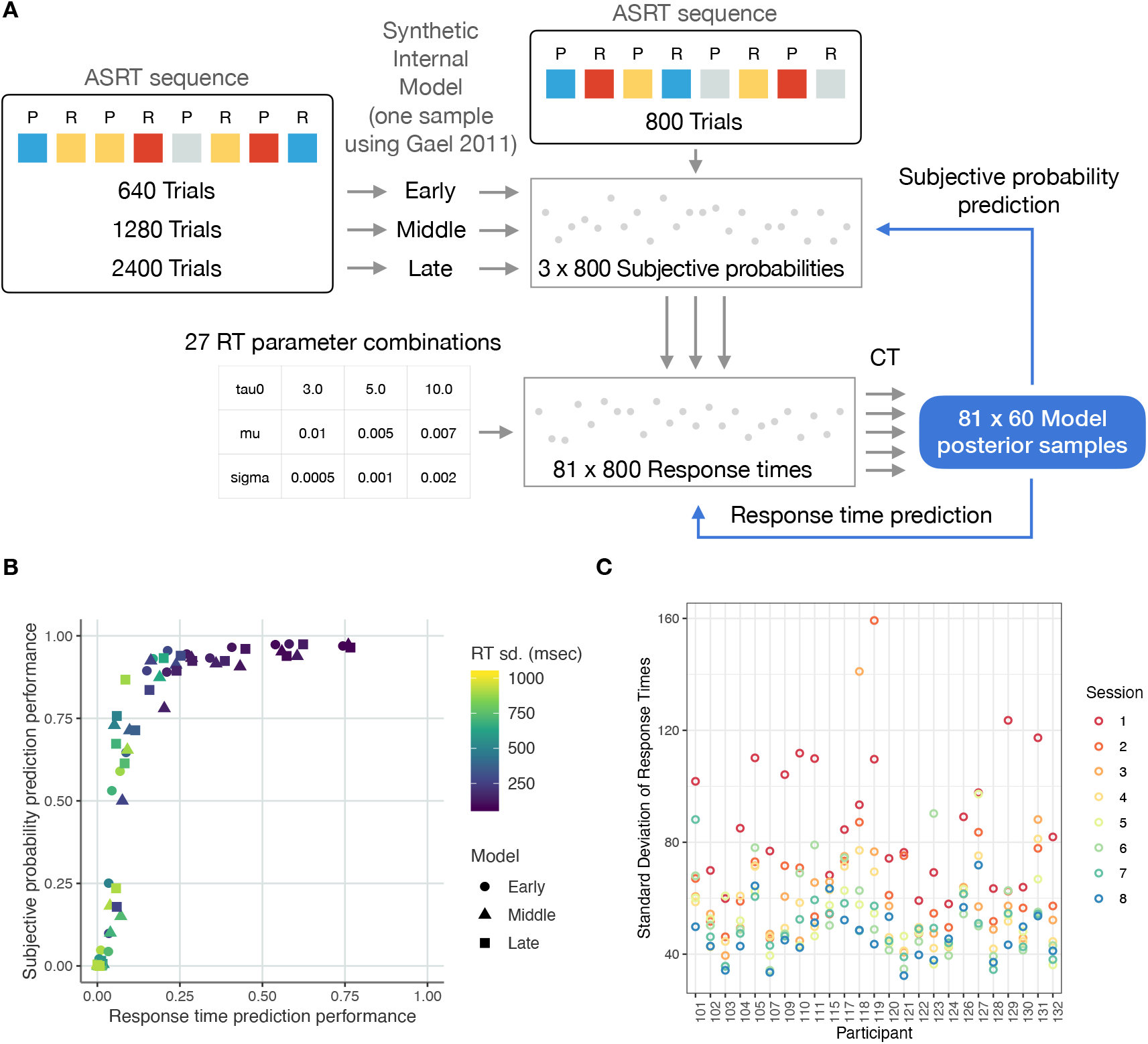
Synthetic data experiment. *A* We first sampled three versions of synthetic internal models using the original iHMM inference method in ***Gael et al. 2008***. The internal models of the synthetic participants differ in their experience (as how many ASRT trials they had seen) - resulting in an “early”, “middle” and “late” model. Then, we generated subjective probability values for each model on a new set of ASRT stimuli (holding the pattern sequence intact). We ran our inference method for 81 synthetic datasets with different parameter settings (*symbols with different colors and shapes*). We use the same number of response times as with the human participants to recover the internal models. *B* Results of our synthetic data experiment. *Symbol colours* correspond to the response time standard deviation. The result shows that while the response time prediction may be at a lower level, the latent predictive probabilities can still be inferred with relatively high accuracy. This shows the inference method can recover the latent structure from a generated response time sequence. *C* Standard deviations of response times of individuals in the first eight experimental sessions.

#### Internal model of participants

The model, established in prior literature, sorts trials into High probability and Low probability triplets. This is equivalent to assuming that the participant uses a two-back (or trigram) model for prediction, predicting the most-likely stimulus conditioning on the previous two observations. Due to the ground truth generative model of the task there is no practical dependence on the identity of the immediately preceding stimulus, and only the penultimate stimulus can contribute to making predictions.

#### Inference

The trigram model has no parameters fitted. Performance is evaluated by the *R*^2^ measure between the response times and the binary variable (high vs low trials) provided by the trigram model.

### Model Comparison

Since not all models considered are Bayesian (i.e. provide an explicit marginal log-likelihood for the response times), we chose to compare models based on explained variance of response times on a test set. Each model produces response time predictions for each trial and each individual separately. When evaluating on a given test set, in order to control for a shift in mean not related to the inherent structure of the response times, we use *R*^2^ as our performance metric. That is equivalent to assuming that the actual observed response times come from a linear model with the predicted response time as mean and an additive homoscedastic (equal variance irrespective of predicted response time) normal noise term.

The *R*^2^ values were calculated separately for each individual’s trials.

### Train and Test Datasets

For the reason described in the above paragraph, for each day (out of 10) of the experiment, out of the 25 blocks each day, we selected blocks 11-20 as a training dataset and blocks 1-10 as test datasets. The main reason for this choice is that on each day in the initial few blocks participants may be engaged in a warm-up phenomenon which fundamentally alters their behaviour in the task. If we use the first 10 blocks as test data, the performance metric may be influenced by 10-30% depending on how many blocks include altered behaviour. However, if we used this part as training data, the whole internal model inference would shift fundamentally, since our inference algorithm assumes a fixed model the entirety of the 10 blocks.

During model inference (train dataset) and performance evaluation (test dataset) the first five random trials and all incorrect response trials’ response times are not considered.

## Statistical Methods

Normality was not checked prior to t-test comparisons. All reported correlations were computed using Pearson’s correlation. T-tests are paired sample tests whenever there is a within-subject comparison. All binomial tests are one-sided. For effect sizes we calculated Cohen’s d using the lsr R package.

### Error prediction

Just as with response time prediction, the model outputs (for each posterior model sample) a subjective probability estimate for each one of the four possible stimuli for each trial. Then, we take mean over these probability estimates over last 60 unique samples of each chain. We decided on using 60 samples since model performances saturate at this number. On Fig. 5 we compute the rank among the four probability estimates of the stimulus and the choice in correct and incorrect trials. Then, based on the subjective probability estimates of the actual occurring stimulus, we plot the receiver operating characteristic curve for predicting whether a given trial will result in an error. This is done by moving a threshold value from 0 to 1 and predicting correct trial if the subjective probability of the upcoming stimulus is above the threshold and an erroneous trial otherwise. The trigram model has two points (other than the (0, 0) and (1, 1) points). This is because the trigram model predicts 0.25 probability for the all stimuli for the first two trials in each block and 0.625 probability for the more high probability trigram element in all other trials and 0.125 for the other stimuli.

## Supplementary Figures

**Figure S4.**
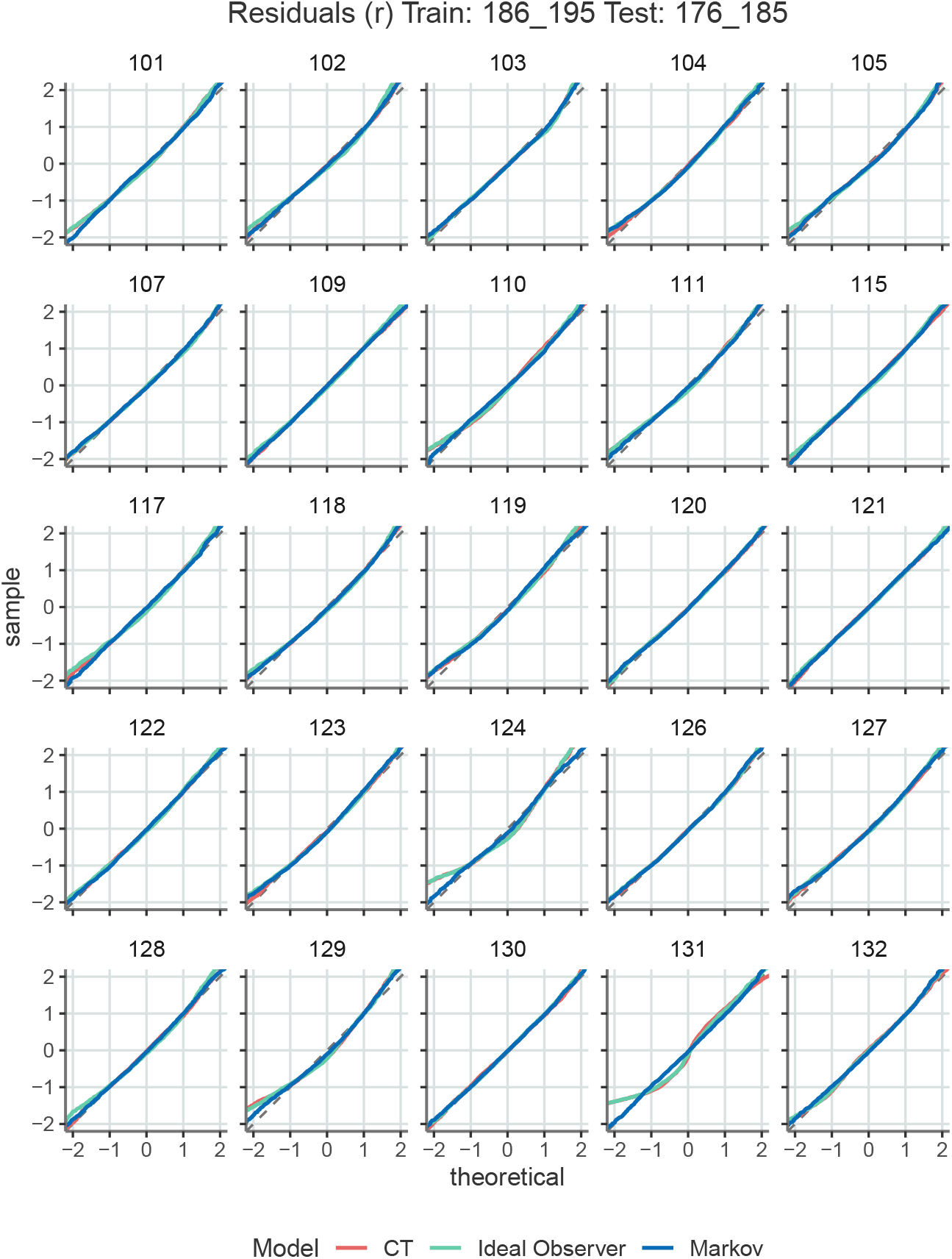
Comparison of quantiles of the (z-scored) *r* variable in the LATER model computed from the response times and predicted subjective probabilities with quantiles of the expected normal distribution for the analysed models (*red*, CT; *green*, ideal observer; *blue*, Markov), also known as QQ-plots. Participant-by-participant shows that the empirical distribution of the *r* parameter on a test set is approximately normal with a few exceptions (see participants 124 131 CT and Ideal Observer models), thus validating model assumptions.

**Figure S5.**
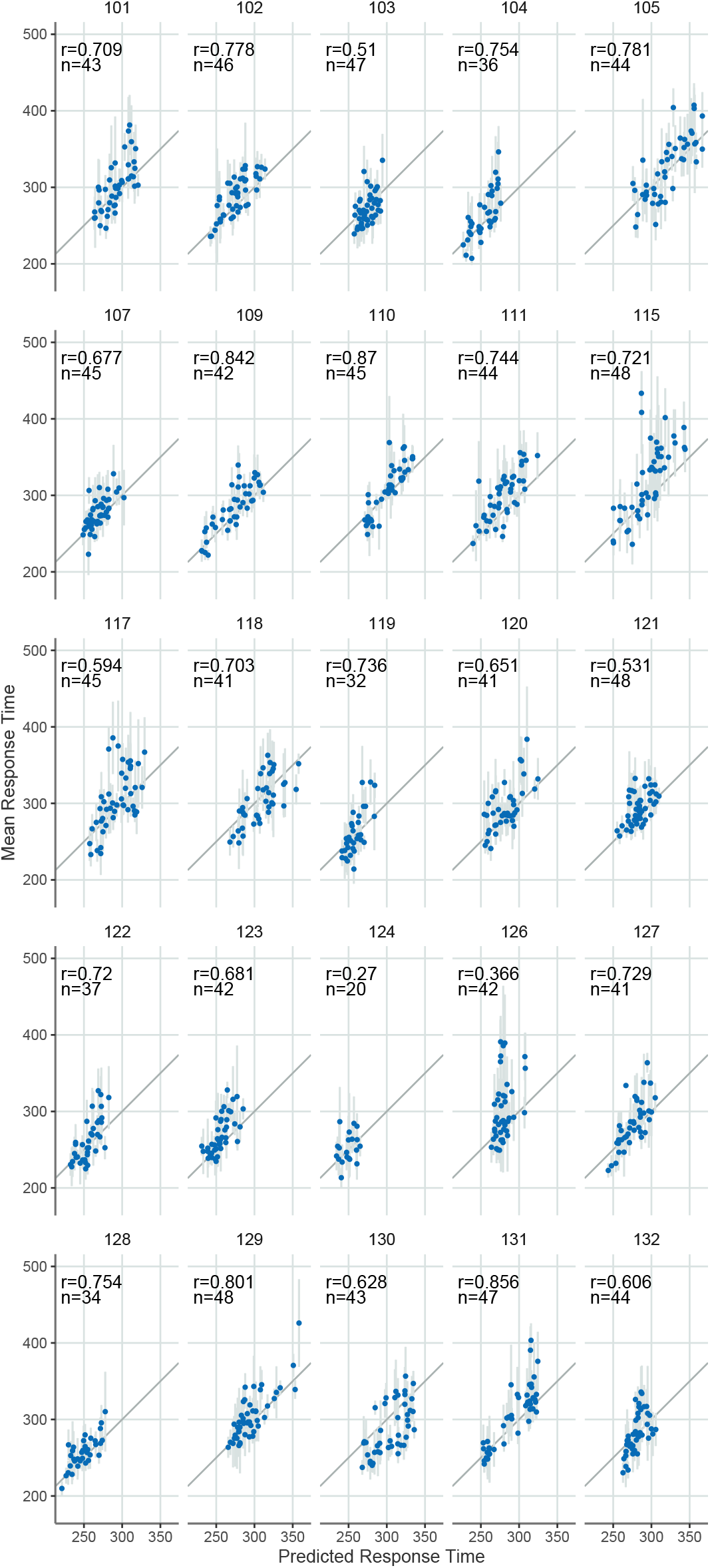
Predicted response time means vs. measured response time means on the test set on Day 8 of the experiment grouped by three element sequences for each participant separately. Only those sequences were included which had at least 5 measured correct response times in order to limit the standard error over the measured response time mean. Error bars show 2 s.e.m.

**Figure S6.**
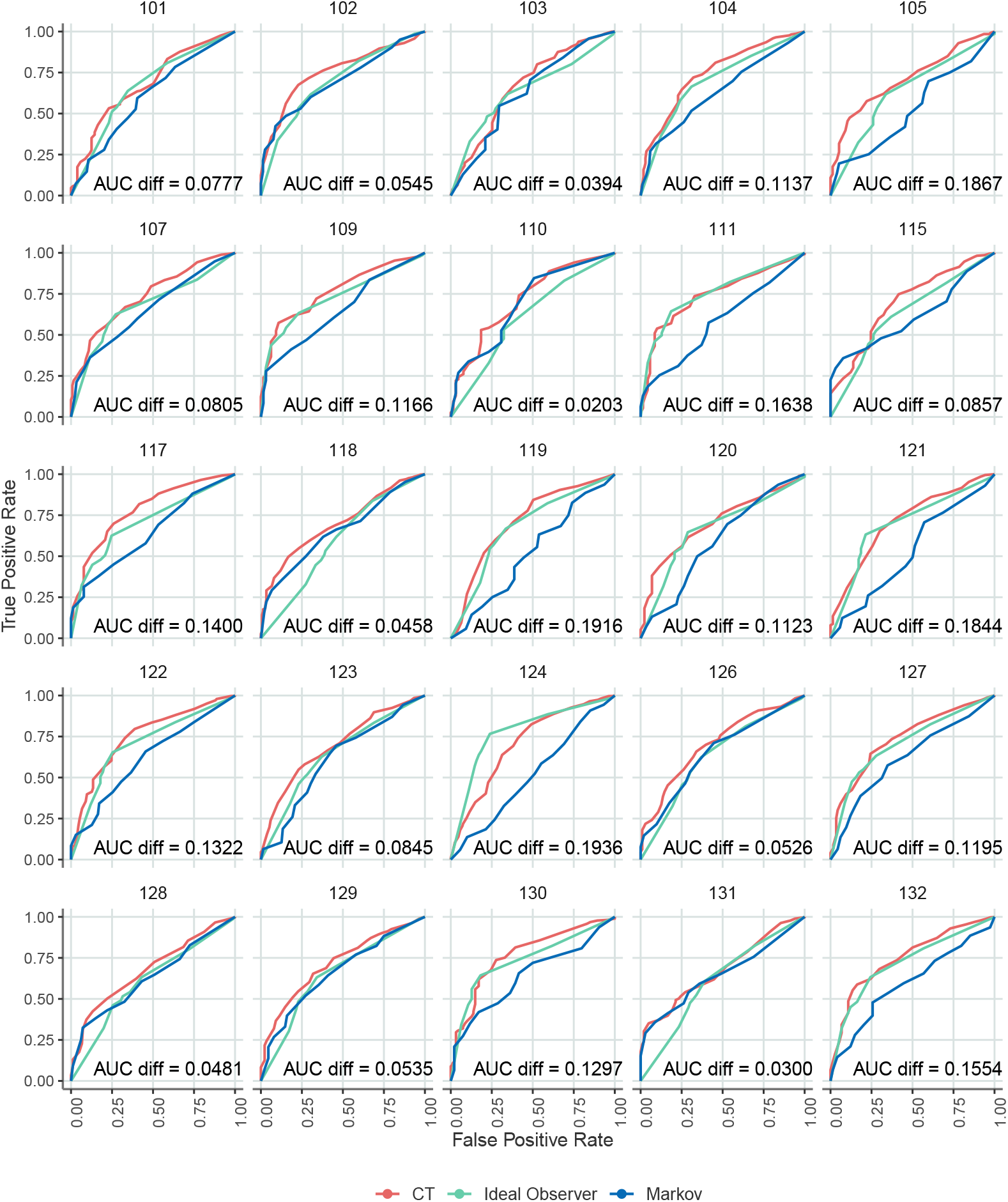
Predicting when errors will occur for each participant individually.

**Figure S7.**
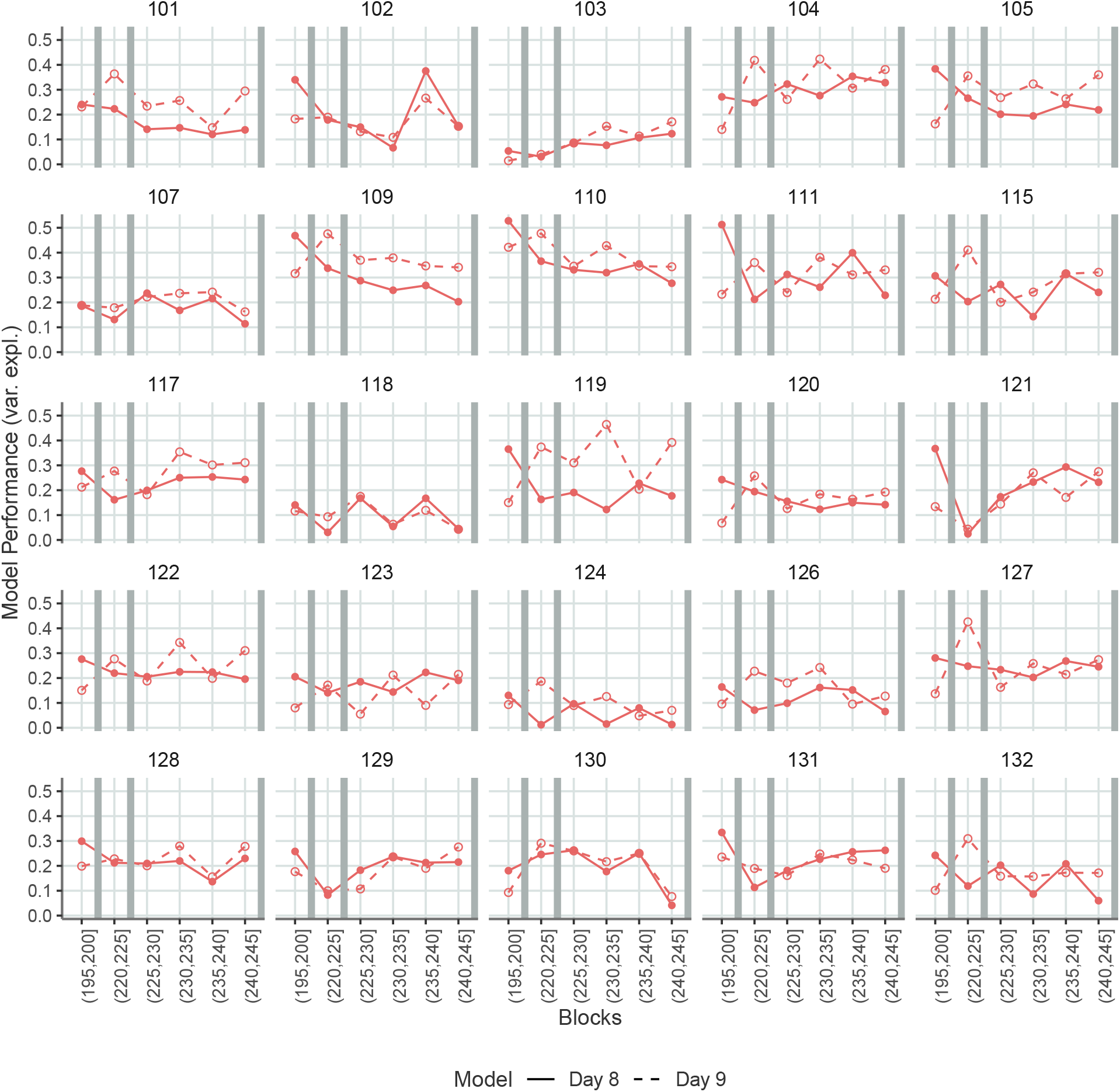
Model performances of CT models trained on Day 8 and Day 9 for each individual.

**Figure S8.**
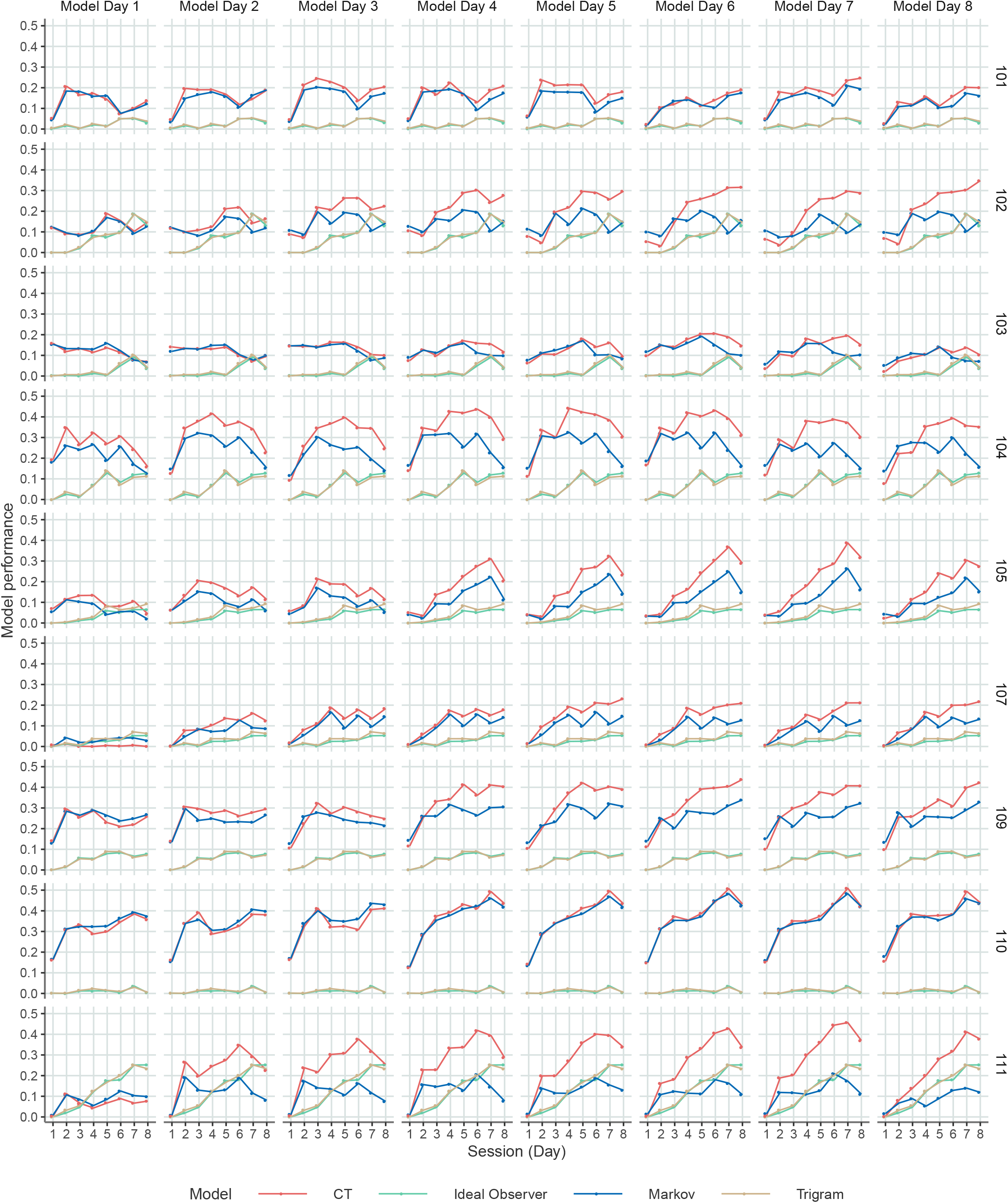
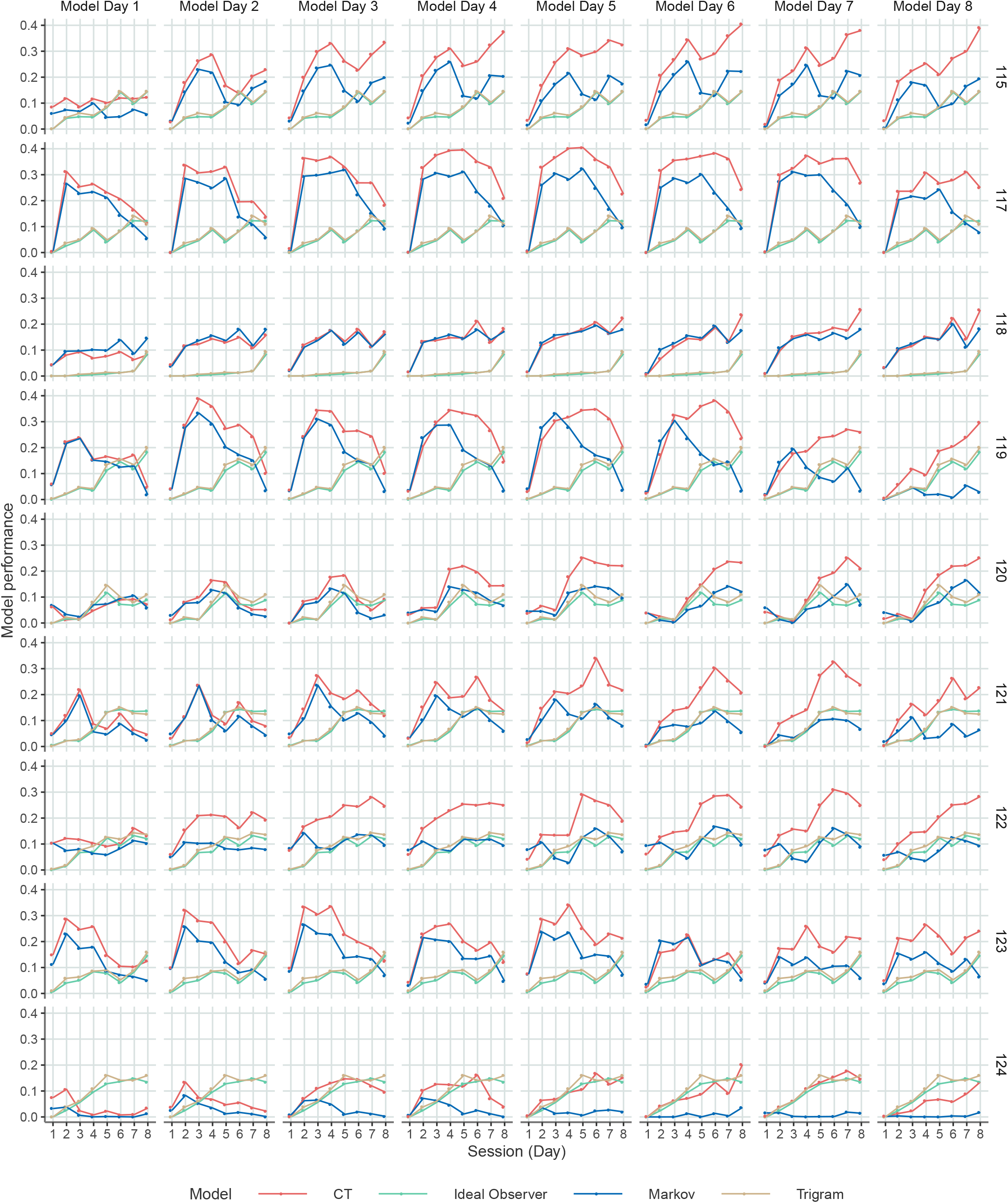
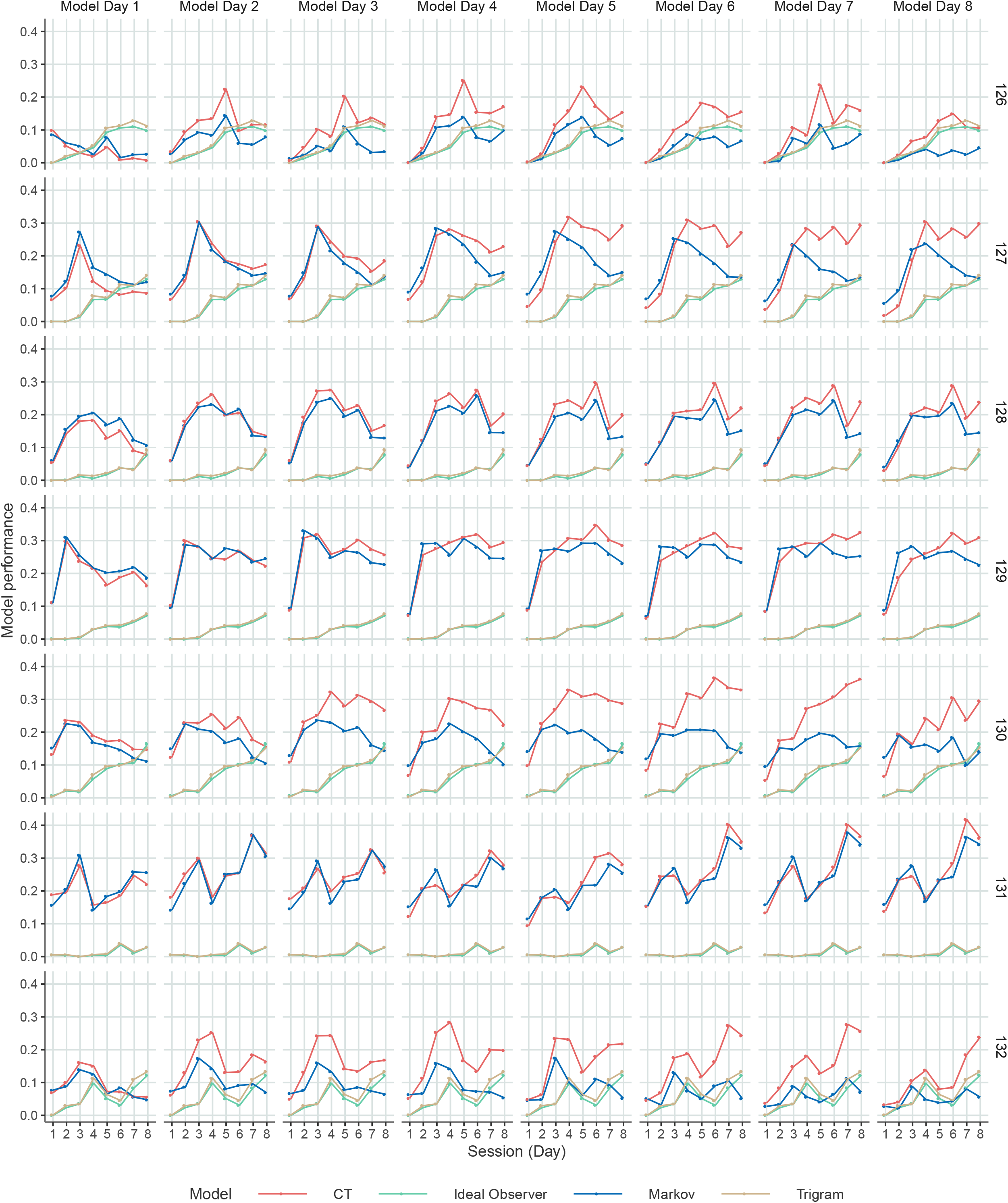
Performances of models trained on different days for each individual. Performances are calculated with RTs above mean + 3.s.d. cut off. This is done to decrease variance of performance. For some participants, the CT model and Markov model diverges, which indicates these individuals infer some non-Markovian structure.

---

**Figure S9.**
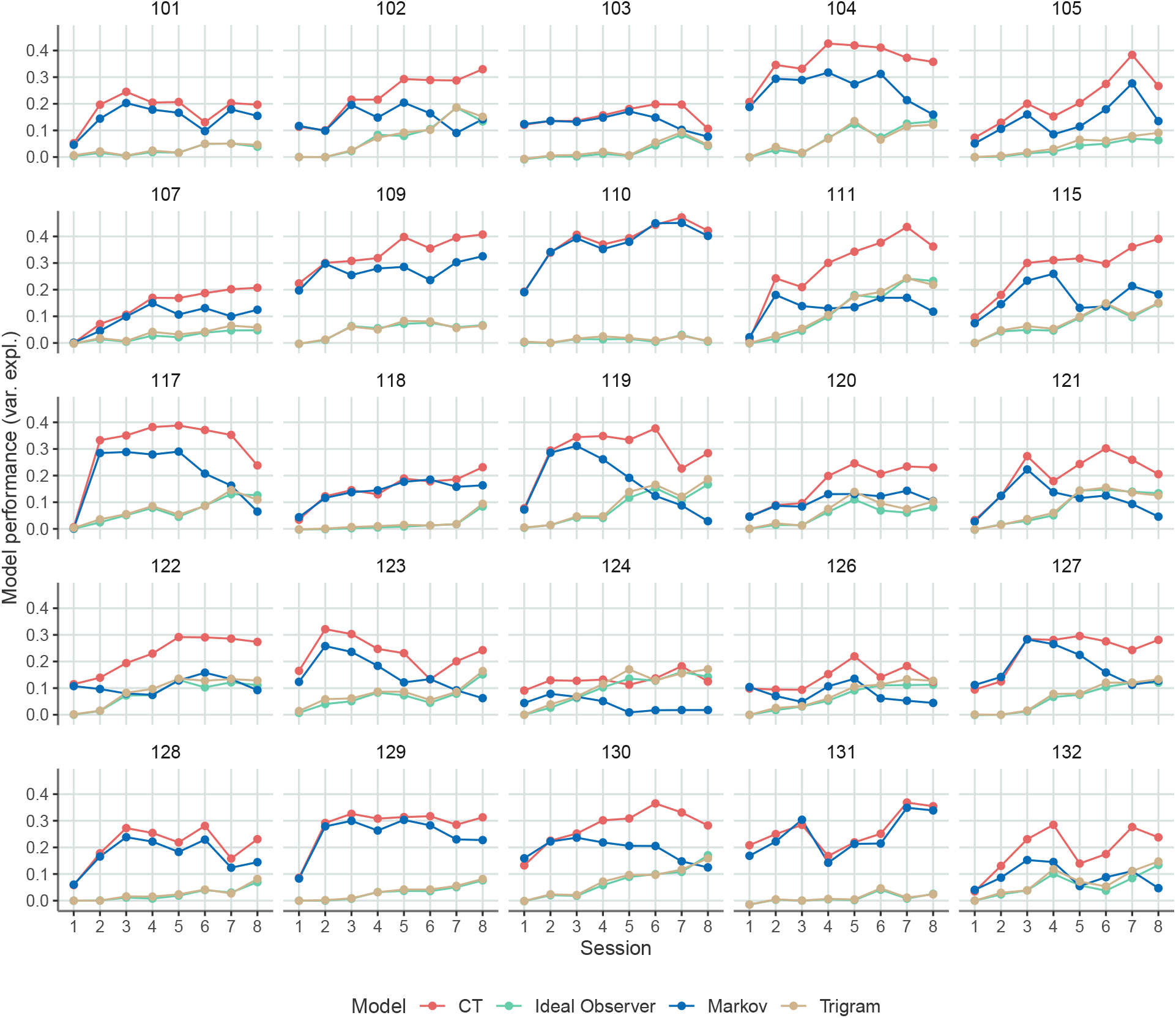
Model performances for all models and all participants individually.

**Figure S10.**
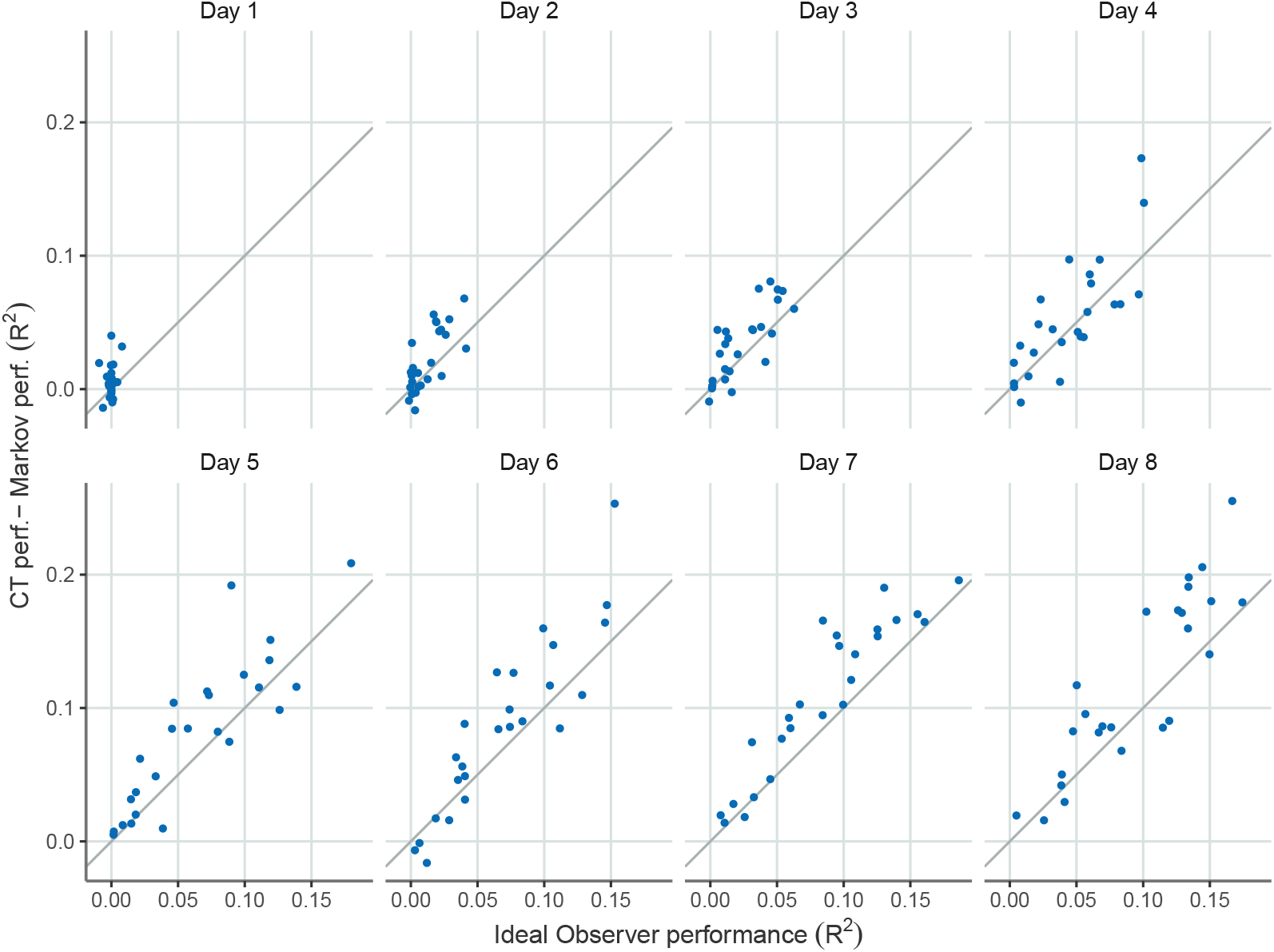
Normalized CT performance as a function of the ideal observer model performance on different days of the experiment. Dots indicate the performance of the models for different individuals.

## Notes

### Competing Interest Statement

The authors have declared no competing interest.

### Summary of Updates

Corrected typo in abstract.

